# Rev-erbα inhibits vascular calcification by repressing osteogenic reprogramming in atherosclerosis

**DOI:** 10.64898/2026.10.14.758688

**Authors:** L Ferri, C Bellengier, M Diedisheim, A Bongiovanni, JB Julla, S Delhaye, GH Kara-Ali, C Duhem, S Farge, Q Thorel, A Hebras, M Leduc, S Quemener, D Eberlé, D Dombrowicz, A Dupont, S Susen, M Rosa, S Germain, Y Sebti, M Tardivel, N Venteclef, JT Haas, B Staels, JF Gauthier, H Duez, B Pourcet

**Affiliations:** Univ. Lille, INSERM, CHU Lille, Institut Pasteur de Lille, U1011-EGID, Lille, France; INEM Institut Necker-Enfants Malades, INSERM U1151, CNRS UMR 8253, Universite Paris Cite, Paris, France; Clinique Saint Gatien Alliance (NCT+), Saint-Cyr-sur-Loire, France; Univ. Lille, CNRS, Inserm, CHU Lille, Institut Pasteur de Lille, BioImaging Center Lille US 41 - UAR 2014 - PLBS, F-59000 Lille, France; INSERM-U955, Univ. Paris Est Créteil, IMRB, Créteil, F-94010, France

**Keywords:** Vascular Calcification, Smooth Muscle Cell, Diabetes, Atherosclerosis, Nuclear Receptors, Rev-erb

## Abstract

**Background:** Atherosclerotic plaque rupture is the most deleterious atherogenic event, responsible for the majority of fatal acute atherothrombotic and myocardial infarctions. Intimal microcalcification is a major cause of atherosclerotic plaque instability. Yet, a thorough characterization of the mechanisms controlling intimal calcification is needed to develop innovative therapeutic strategies to treat high risk patients. The nuclear receptor Rev-erbα regulates lipoprotein metabolism and exerts anti-inflammatory activities. Here, we hypothesized that Rev-erbα modulates atherogenesis by regulating arterial calcification processes.

**Methods:** We used a multi-omics approach in the ANGIOSAFE and the ATHEROPORTAL human cohort to identify pathways and factors associated with vascular calcification in diabetic patients. We then compared *REV-ERBα* expression between calcified and non-calcified areas of human endarterectomies using laser capture microdissection assays (n=11-22/group). We further assessed the functional role of Rev-erbα on calcium deposition and expression of osteoblast markers in aortic roots of 18-month-old *Ldlr*^*–/–*^*Rev-erbα*^*–/–*^ and *Ldlr*^*–/–*^*Rev-erbα*^*+/+*^ mice (n=10/group). *Rev-erbα*-dependent molecular pathways that drive calcification were identified using gene expression profiling in aortic plaques from *Ldlr*^*–/–*^*Rev-erbα*^*–/–*^ and *Ldlr*^*–/–*^*Rev-erbα*^*+/+*^ mice (n=8–10/group). We then confirmed omics data *in vitro* using primary aortic smooth muscle cells (SMCs) (n=3-6) and assessing inflammation-driven calcification process as well as Rev-erbα-dependent transcriptional mechanisms.

**Results:** In the ANGIOSAFE cohort (n=668), *REV-ERBα* expression is significantly lower in circulating classical CD14^+^CD16^-^ monocytes from diabetic patients with low coronary artery calcium (CAC) compared to those exhibiting high CAC scores. In human atheromatous lesions from the AtheroPortal cohort (N=968), *REV-ERBα* expression is lower in highly calcified plaques from patients with diabetes mellitus compared to those with no diabetes or no calcification. Consistently, in laser capture microdissection experiments, *REV-ERBα* expression is markedly lower in calcified areas of human endarterectomy samples, compared to non-calcified regions. *Rev-erbα* deficient 18-months-old *Ldlr^−/−^* mice exhibit accelerated atherogenesis and vascular calcification, and aortic smooth muscle cells (SMCs) exhibit an extensive reprogramming toward an osteochondroblastic phenotype. Functional *in vitro* studies show that Rev-erbα maintains SMC identity by preventing their IL-1β-driven transdifferentiation into osteochondroblast-like cells, notably through the repression of key osteogenic regulators, including *Runx2, Spp1, Osterix* and *Sox9*. Finally, we demonstrate that Rev-erbα represses the hyaluronidase Cemip expression during SMC transdifferentiation. Because hyaluronic acid degradation promotes intimal calcification, we provide here a mechanistic link between Rev-erbα activity, extracellular matrix remodelling, and vascular calcification.

**Conclusions:** Rev-erbα is identified as an important brake on SMC-driven vascular calcification, suppressing both osteogenic reprogramming and matrix-remodeling pathways. Thus, this study opens new avenues for developing novel therapeutic tools targeting Rev-erbα to reduce vascular calcification and associated cardiovascular risk in high-risk patients.

## Introduction

Cardiovascular diseases account for 35% of global mortality, representing the leading cause of death worldwide^1^. Importantly, 84% of cardiovascular-related deaths are due to acute events, *e*.*g*. ischemic heart diseases and stroke, that are largely driven by atherosclerotic plaque instability and rupture^2^. Despite its clinical significance, the cellular and molecular mechanisms underlying plaque rupture are still poorly understood.

Atherosclerosis is a chronic inflammatory disease driven by lipid accumulation in the subendothelial space, which triggers inflammation, leukocyte infiltration and smooth muscle cell (SMC) activation^3,4^. Vascular calcification plays a central role in plaque heterogeneity, instability and rupture^5^. In clinical practice, the coronary artery calcium (CAC) score, reflecting vascular calcification, is therefore commonly used as a surrogate of cardiovascular risk and plaque instability^6^.

Vascular calcification consists in the progressive deposition of calcium crystals affecting nearly all arterial beds, within both the intimal and medial layers. Intimal microcalcifications (<15 µm), the most frequent and clinically relevant form of vascular calcification, correlate positively with atherogenesis and plaque rupture^5^, whilst macrocalcifications (>15 µm) increase arterial stiffness in patients with chronic kidney disease^7^. While macrocalcifications are generally considered to stabilize atherosclerotic plaques, they however originate from the aggregation of spotty microcalcifications, whose shape and protrusion into the fibrous cap contribute to plaque instability and rupture^8,9^.

Smooth muscle cells (SMCs) play a key detrimental role in this process, which is amplified by the inflammatory context of the atherosclerotic plaque. Indeed, SMCs display a high level of plasticity^10^ since they can transdifferentiate into mesenchymal stem cells, fibroblast-like cells, foam cells, macrophage-like cells or osteogenic-like cells^11–13^. SMCs initially undergo a phenotypic switch from a contractile to a secretory phenotype in response to calcium and phosphate released by necroptotic macrophages and SMCs^14^. Lesional macrophages further amplify this process by secreting pro-inflammatory cytokines, notably IL-1β, which promotes the transition of secretory SMCs into osteochondroblast-like cells, thereby enhancing extracellular matrix mineralization^15^. At the molecular level, this osteogenic reprogramming is characterized by the induction of transcription factors such as *Sox9, Runx2* and *osterix/Sp7* in response to pro-calcifying and pro-inflammatory conditions^14^. The Sox9, Runx2 and osterix transcription factors control the expression of functional pro-calcifying factors including osteocalcin, whose expression is increased in calcified aortas and controls vascular calcification^16,17^, the tissue non-specific alkaline phosphatase (TNAP), osteopontin (Spp1), RANK ligand (RANKL), extracellular matrix components such as type I collagen or anti-calcifying proteins such as Matrix Gla Protein (MGP)^14^. TNAP converts the inorganic pyrophosphate (PPi) into inorganic phosphate (Pi) to form hydroxyapatite crystals, whose anchorage to type I collagen fibers is stimulated by osteocalcin^18^. ECM composition and remodeling, together with cell proliferation, migration and differentiation are determinants of plaque stability. Moreover, diabetic patients display an even greater complexity of atherosclerotic plaques compared with non-diabetic individuals, which has been associated with dysregulated vascular wall hyaluronic acid content, leading to impaired ECM remodeling^19^. The hyaluronidase Cell migration inducing protein (Cemip), which depolymerizes hyaluronic acid into small fragments, controls Wnt/beta-catenin-dependent proliferation and migration of SMCs in atherosclerotic plaques^20^. Degradation of the key ECM component hyaluronic acid into small fragments promotes vascular calcification^21,22^. Altered ECM regulation thus likely contributes to plaque destabilization and to the initiation of ectopic calcification.

In this study, we aim to identify key factors that restrain vascular calcification in humans. We identify *REV-ERBα* as an important factor associated with vascular calcification in plaque from diabetic patients at high cardiovascular risk. REV-ERBα is a transcriptional repressor belonging to the nuclear receptor superfamily, which regulates metabolism, inflammation and circadian rhythmicity^23–25^ and whose activity can be modulated by natural ligands such as hemin, or synthetic ligands such as SR9009^23,26^. Notably, *REV-ERBα* expression is reduced in plaque from diabetic patients with severe calcification and in calcified versus non-calcified regions of human endarterectomy biopsies. By combining *in vivo* mouse model, single-cell transcriptomic and primary aortic SMC cultures, we further show that Rev-erbα protects from vascular calcification by repressing IL-1β-driven osteochondrogenic reprogramming of SMCs and inhibiting the hyaluronidase Cemip. Altogether, these findings identify Rev-erbα as a key regulator of atherosclerotic plaque progression and calcification.

## Materials and Methods

### Human cohorts

#### ANGIOSAFE cohort

AngioSafe-2 cohort was registered at ClinicalTrials.gov (Identifier: NCT02671864). A total of 668 participants with type 2 diabetes (T2D) were stratified according to their cardiovascular risk using the Coronary Artery Calcium (CAC) score: low CAC (Q1, n=218), moderate CAC (Q2, n=220; Q3, n=128), and high CAC (Q4, n=102). The study was approved by the relevant institutional review boards and ethics committees, including the CPP Île-de-France Ethics Committee, which authorized the inclusion of all individuals.

#### OPAL Lille cohort

Human atherosclerotic plaques were removed from patients eligible for surgical carotid endarterectomy (significant carotid artery stenosis >70%) recruited at the Cardiovascular Surgery Department, Lille University Hospital, France27. The study protocol was approved by the local ethics committee (OPAL-Lille study, CPP11-15 n°2010-A01401-38) in accordance with the ethical guidelines of the Declaration of Helsinki. Written informed consent was obtained from all subjects.

#### AtheroPortal

Bulk RNA-sequencing data were analysed from the Athero-Express Biobank Study, an ongoing prospective longitudinal cohort of patients undergoing carotid or femoral endarterectomy at two tertiary centers in the Netherlands. The study design has been described previously^28^. Transcriptomic data from 1,093 atherosclerotic plaque segments (AERNAS1, *n*=622; AERNAS2, *n*=471) together with the corresponding clinical and plaque phenotyping data were accessed through the AtheroPortal (Peters TS, Krebbers S, van der Laan SW, *et al*., *medRxiv*, 2025; accessed on june 2026 at https://atheroportal.nl/app/). A Kruskal-Wallis test was used to determine whether there are statistically significant differences between groups. Comparison between groups is performed using t-test for each group pair, and p-values are corrected for multiple testing using the Bonferroni correction.

### Statistical Analysis

Statistical analysis was performed on GraphPad Prism software (version 11.0.2) using either *t* test after the normality is verified or Mann-Whitney test for comparisons between 2 groups and one-way and Kruskal-Wallis or two-way ANOVA, after the normality is verified, with Šídák or *post-hoc* multiple comparisons of more than 2 groups, as indicated in the legends. An identification of statistical outliers was performed with a ROUT (Q=1%). Differences were considered significant when *p*<0.05. All *in vitro* experiments were repeated independently at least three times, and representative results are presented.

A detailed description of all methods is available in the Supplemental Material.

## Results

### REV-ERBα is associated with plaque complexity in diabetic patients and with vascular calcification in human endarterectomies

To investigate the mechanisms driving plaque complexity and intimal calcification, as well as identify novel contributing factors, we first studied diabetic patients, who are at increased cardiovascular risk and tend to have more complex, calcified plaques than non-diabetic individuals^36^. Given that systemic inflammation and the number of circulating immune cells are a good proxy for cardiovascular risk^37^, we first analyzed the peripheral blood cells from diabetic patients enrolled in the ANGIOSAFE cohort^38^. Diabetic patients were stratified into 4 cardiovascular risk quartiles based on their CAC score (Figure 1A) where a CAC score >300 is associated with a higher risk of cardiovascular events and mortality^39^. Compared with patients in the first quartile, corresponding to the lowest cardiovascular risk (Q1, with a CAC score <1), those in the fourth quartile (Q4, with a CAC score >400) displayed elevated blood total leukocytes, and in particular monocytes (Suppl. Figure 1A-B)^37,38,40^. Indeed, the frequency of classical CD14^+^CD16^-^ monocytes increases (Supplementary figure 1C), whereas the percentage of non-classical CD14^low^CD16^+^ monocytes is lower in Q3 compared to Q1 patients (Supplementary figure 1D)^41^. We then assessed differences in classical CD14^+^CD16^-^ monocytes between Q1 and Q4 patients by identifying differentially regulated genes using bulk RNA sequencing. Genes upregulated in Q4 patients, compared to Q1 patients, were mostly involved in “DNA modification” biological processes highlighting a substantial transcriptional reprogramming of these cells (Figure 1B). Precisely, GO molecular pathway analysis indicates that pathways most profoundly different between Q1 and Q4 patients are primarily associated with transcriptional repression, including histone deacetylase activity, protein lysine deacetylase activity and histone methyltransferase binding (Figure 1C). Among transcription factors that are differentially regulated between Q1 and Q4 classical CD14^+^CD16^-^ monocytes, we found the nuclear receptor REV-ERBα (alias NR1D1) as the only repressor known to regulate both inflammation^24^ and metabolism^42^ (Figure 1D). As expected, *REV-ERBα* expression is induced in anti-inflammatory classical CD14^+^CD16^-^ monocytes isolated from high (Q4) compared to low (Q1) cardiovascular risk patients (Figure 1D, Supplementary Figure 1E). On the other hand, the expression of *REV-ERBα* is significantly lower in highly calcified atheromatous compared to non-calcified plaques from diabetic patients in the AtheroPortal human cohort (Supplementary Figure 1F) (Peters TS, Krebbers S, van der Laan SW, *et al*., *medRxiv*, 2025; accessed on june 2026 at (https://atheroportal.nl/app/). These observations raise the hypothesis for a protective role of REV-ERBα against vascular calcification in atherosclerotic plaque.

**Figure 1:**
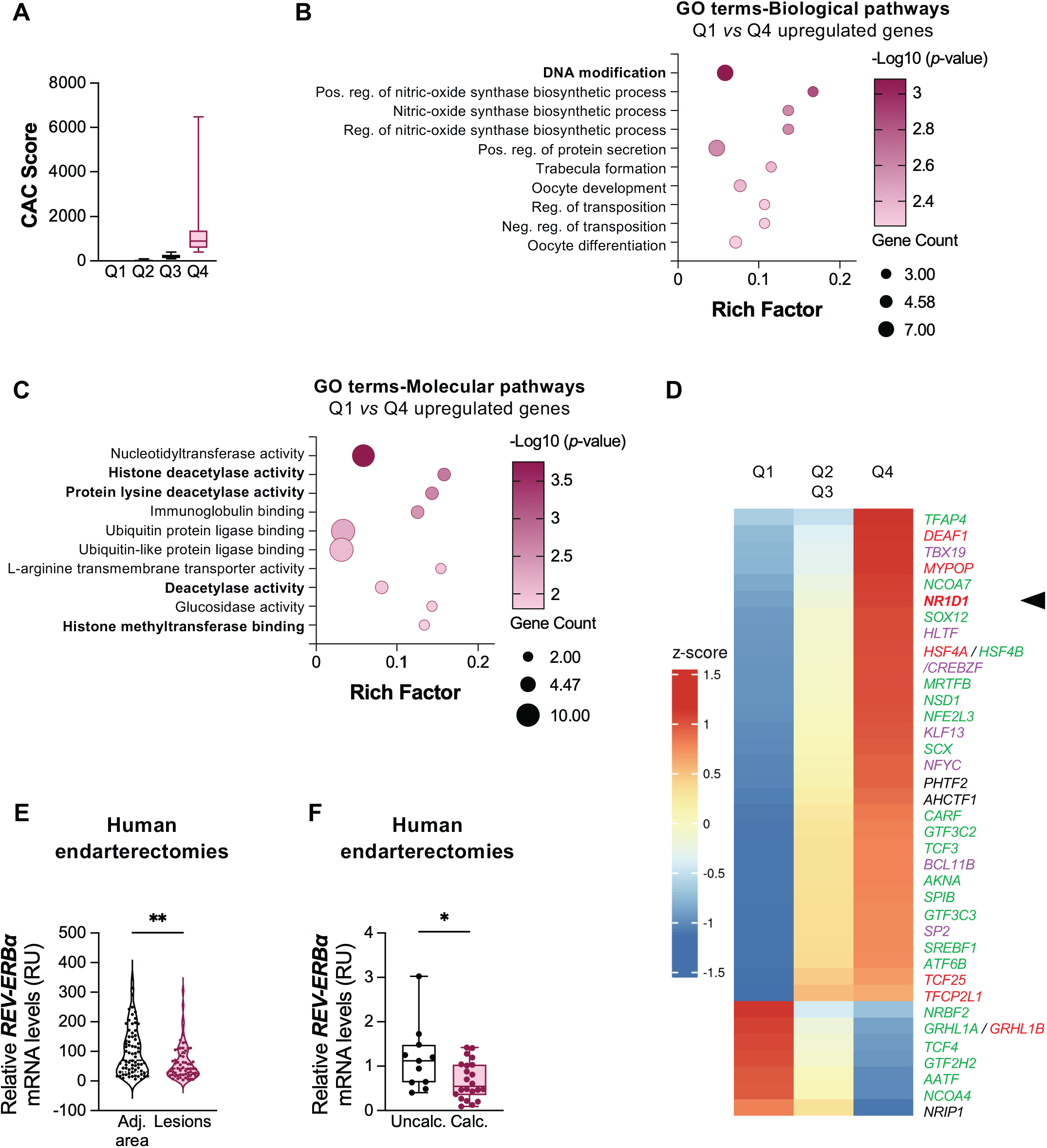
Gene expression profile of diabetic patients stratified by calcium score quartile and analysis of REV-ERBα expression in distinct regions of atherosclerotic plaques. **A**. Quartile distribution of diabetic patients according to their CAC score, classified as low, intermediate, and high (Q1: n=218, Q2: n=220, Q3: n=128, Q4: n=102). Data were analyzed using a one-way ordinary ANOVA test. **B-D**. Representative gene ontology (GO) enrichment analyses of biological processes (B), molecular pathways (C) and heat map illustrating the expression levels of transcription factors (D) based on RNA-seq data from CD14^+^CD16^-^ monocytes isolated from blood samples of Angiosafe diabetic patients classified into quartiles according to their CAC score: low (Q1), moderate (Q2-Q3), and high (Q4) (n=11/group). Transcription factor names are color-coded according to their known transcriptional activity : green, transcriptional activators; red, transcriptional repressors; purple, both activators and repressors; and black, DNA-binding transcription factors for which the transcriptional activity (activator or repressor) remains unknown. **E-F**. Relative levels of REV-ERBα mRNA in human endarterectomy samples from OPAL cohorts. Relative REV-ERBα mRNA expression in atherogenic (n=64) versus healthy regions (n=72) of human endarterectomies from the OPAL cohort (E) (violin plots; Mann-Whitney test, ** *p*<0.01) and in non-calcified (n = 11) *versus* calcified (n = 22) areas from the OPAL cohort (F) (mean ± SD; Mann-Whitney test, \**p < 0*.*05*).

We next compared the expression of *REV-ERBα* in atherosclerotic lesions obtained from the OPAL human endarterectomy biobank^43^ using laser microdissection analysis. *REV-ERBα* expression is lower in atherosclerotic lesion areas compared to adjacent healthy areas, again suggesting a protective role against atherogenesis (Figure 1E). Strikingly, lesion *REV-ERBα* expression levels are higher in non-calcified compared to calcified areas, evoking an anti-calcifying role (Figure 1F, Supplementary Figure 1H). It is noteworthy that REV-ERBβ, an isoform displaying redundant activities with REV-ERBα, is not differentially expressed between calcified and non-calcified regions (Supplementary Figure 1G), indicating a specific role for REV-ERBα in plaque calcification. Altogether, our data associate low intra-plaque *REV-ERBα* expression levels with atherogenesis and vascular calcification in humans.

### Rev-erbα controls atherogenesis and vascular calcification *in vivo*

To determine whether Rev-erbα contributes to the development of complex atherosclerotic plaques or whether its expression only reflects plaque heterogeneity, we assessed atherosclerotic plaque development in aged 18-month-old *Ldlr*^−/−^ littermate male mice deficient or not for *Rev-erbα* (*Ldlr*^−/−^*Rev-erbα*^*+/+*^*vs. Ldlr*^−/−^*Rev-erbα*^−/−^) and fed a chow diet. The *Ldlr*^−/−^ mouse model progressively develops complex atherosclerotic lesions with calcified plaques, thereby closely modeling the human pathogenesis^44^. Interestingly, *Ldlr*^−/−^*Rev-erbα*^−/−^ mice display larger lesions in aortic roots compared to littermate controls (Figure 2A-C), accompanied by increased collagen deposition, macrophages and CD4^+^ T cell surface areas (Figure 2D-F). However, when normalized to plaque area, collagen and leukocyte surface areas were not different between *Ldlr*^−/−^*Rev-erbα*^*+/+*^ and *Ldlr*^−/−^*Rev-erbα*^−/−^ mice (Figure 2G-I), suggesting that *Rev-erbα* deficiency does neither promote collagen deposition nor monocyte and CD4^+^ T cell recruitment, despite larger plaques in Rev-erbα-*deficient* mice. Since *REV-ERBα* expression is inversely associated with intimal calcification in human plaques (Figure 1F), we next assessed the role of Rev-erbα in vascular calcification by quantifying alizarin red-positive calcified areas in the aortic sinus and the ascending aortas of *Ldlr*^−/−^*Rev-erbα*^*+/+*^ and *Ldlr*^−/−^*Rev-erbα*^−/−^ mice. As expected, calcification occurs in the intimal layer of the vascular wall (Figure 3A) and Rev-erbα deficiency significantly increases calcium deposition both in the aortic sinus (from 0 µm to 1610 µm where 0 µm corresponds to the beginning of aortic valve leaflets) and the ascending aorta (from −2000 µm to 0 µm), even when normalized by the lesion area (Figure 3A-B). Together these data point to a role of Rev-erbα in plaque calcification.

**Figure 2:**
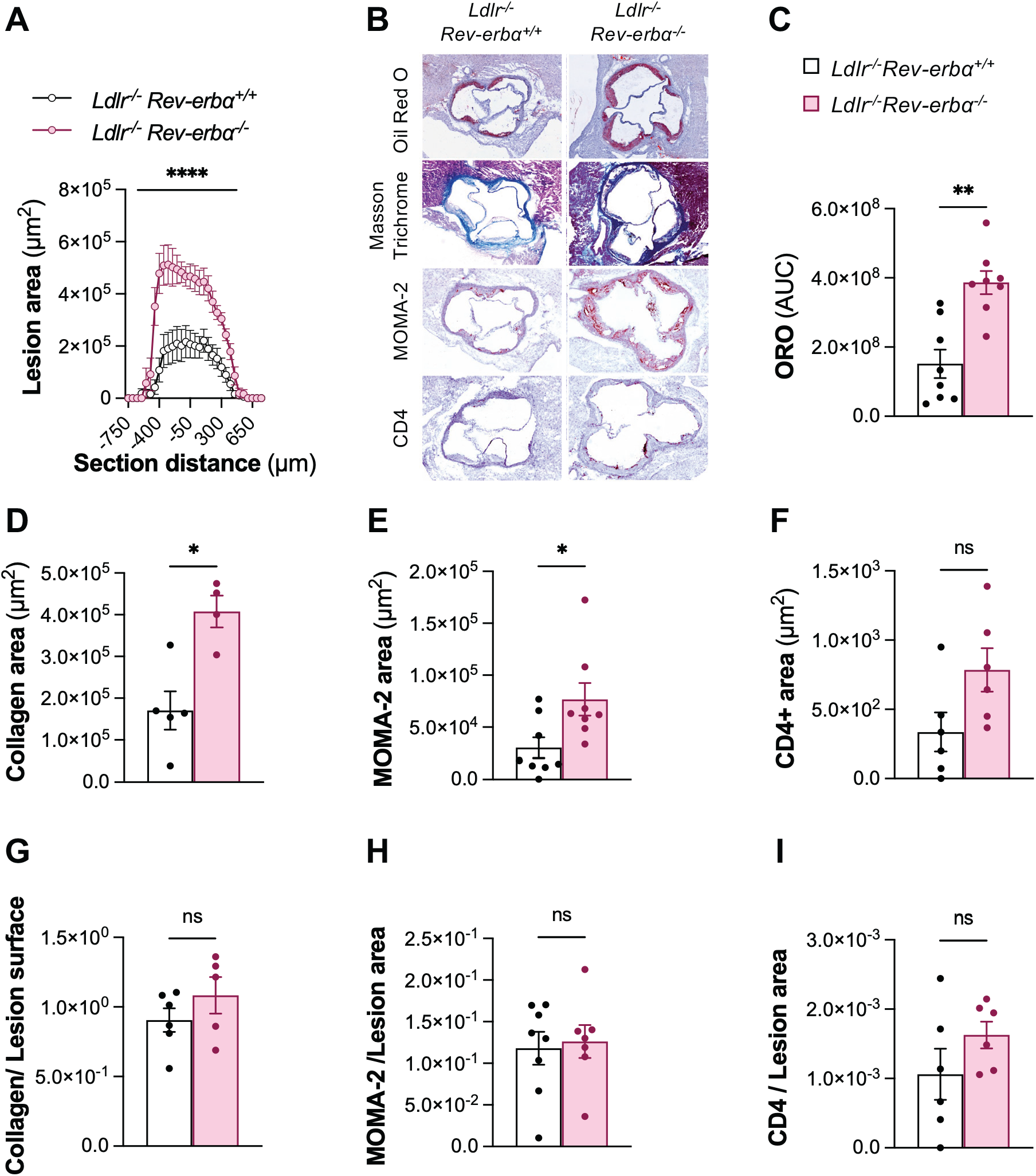
Effect of Rev-erbα deletion on plaque composition in Ldlr^−/−^ mice. **A**. Quantification of atheromatous lesion area in aortic sinus sections from 18-month-old *Ldlr*^−/−^*Rev-erbα*^*+/+*^ and *Ldlr*^−/−^*Rev-erbα*^−/−^ mice, visualized by Oil Red O staining, as a function of distance from the valves. Data are means ± SEM (n=8/genotype), two-way ANOVA, **** *p*<0.0001. **B**. Representative staining for lesion size using Oil Red O, collagen deposition by Trichrome Masson, and macrophage (MOMA-2) and T lymphocyte (CD4^+^) infiltration in aortic sinus sections from 18-month-old *Ldlr*^−/−^*Rev-erbα*^*+/+*^ and *Ldlr*^−/−^*Rev-erbα*^−/−^ mice. Contrast and brightness adjustments were applied equally to each condition using ImageJ2 software (2.16.0/1.5t). **C-I**. Quantification of lipid deposition (C), collagen content (D, G), and macrophage (MOMA-2) (E, H) and T lymphocyte (CD4^+^) (F, I) infiltration either as absolute measures (C–F) or normalized to lesion area (G–I). Data are mean ± SEM (n=8/genotype), Mann-Whitney test, * *p*<0.05, ** *p*<0.01.

**Figure 3:**
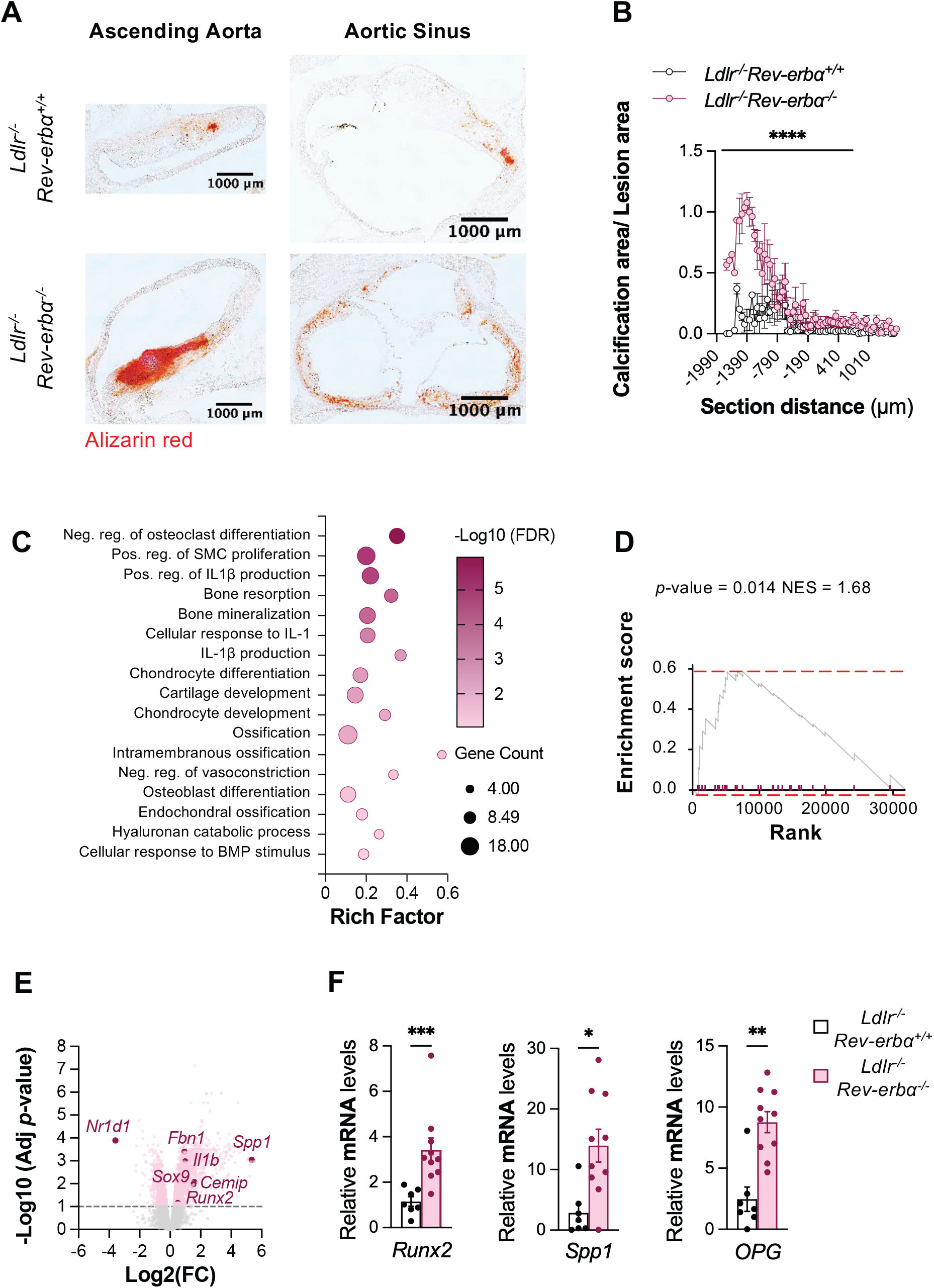
Rev-erb-α play a predominant role in vascular calcification. **A**.Representative calcium deposition staining of ascending aortic and aortic sinus sections from 18-month-old *Ldlr*^−/−^*Rev-erbα*^*+/+*^ and *Ldlr*^−/−^*Rev-erbα*^−/−^ mice after alizarin red staining. Contrast and brightness adjustments were applied equally to each conditions using ImageJ2 software (2.16.0/1.5t).**B**.Quantification of calcium deposits relative to lesion area in ascending aorta and aortic sinus sections after Alizarin Red staining, as a function of distance from the valves. Data are presented as means ± SEM (n=8-10 per genotype). Statistical analysis was performed using a two-way ANOVA test, with significance indicated by **** *p*<0.0001 for AUC. **C**. Enrichment of differentially expressed genes (DEGs) in biological pathways (GO terms BP). **D-E**. Gene set enrichment analysis (GSEA) on genes involved in human vascular calcification using whole aortic DEGs from 18-month-old *Ldlr*^−/−^*Rev-erbα*^*+/+*^ and *Ldlr*^−/−^*Rev-erbα*^−/−^ mice (D), and Volcano plot (E) of whole aortic DEGs. **F**. Osteoblastic marker mRNA levels in whole aortas. Data are shown as means ± SEM (n=7-10 per genotype), Mann-Whitney test, * *p*<0.05, ** *p*<0.01, *** *p*<0.001.

To decipher the molecular and cellular mechanisms by which Rev-erbα controls plaque progression and calcification, we performed a transcriptomic analysis on whole aortas harvested from 12-month-old *Ldlr*^−/−^*Rev-erbα*^*+/+*^ and *Ldlr*^−/−^*Rev-erbα*^−/−^ mice fed a chow diet. As shown in Supplementary Figure 2A-B, the plaque transcriptome of *Ldlr*^−/−^*Rev-erbα*^*+/+*^ mice differs from that of *Ldlr*^−/−^*Rev-erbα*^−/−^ mice. Interestingly, using a GO term enrichment analysis, upregulated genes in *Ldlr*^−/−^*Rev-erbα*^−/−^ mice are enriched in pathways related to inflammation (response and production of IL-1β), SMC proliferation, matrix remodeling (hyaluronan catabolism) as well as bone homeostasis (ossification, bone mineralization, BMP response, osteoblast and chondrocyte differentiation…) (Figure 3C). Accordingly, a fGSEA confirmed that most upregulated genes in *Ldlr*^−/−^*Rev-erbα*^−/−^ belong to inflammatory processes, ECM remodeling and calcifying processes (Supplementary Figure 2C).

Importantly, using a human gene list in a GSEA analysis (Supplementary figure 2D), genes involved in human vascular calcification processes are mostly upregulated in *Ldlr*^−/−^*Rev-erbα*^−/−^ aortas when compared to their littermate controls (Figure 3D). Accordingly, genes controlling osteoblast differentiation, such as *Sox9, Runx2, Spp1* (encoding the osteopontin protein), as well as inflammation (*e*.*g. Il1β*) and extracellular matrix remodeling (*Fbn1, Cemip*), are up-regulated in *Ldlr*^−/−^*Rev-erbα*^−/−^ compared to their littermate control mice (Figure 3E-F). Altogether, our data show that *Rev-erbα* deletion promotes atherosclerosis development with intimal vascular calcification and induces an osteochondroblastic differentiation transcriptional program in the vascular wall during advanced atherogenesis.

### Rev-erbα controls osteoblastic differentiation of smooth muscle cells *in vivo*

To define how Rev-erbα regulates vascular cell plasticity, we performed single-cell RNA sequencing of CD45^−^ cells isolated from atherosclerotic aortas of *Ldlr*^−/−^*Rev-erbα*^*+/+*^ and *Ldlr*^−/−^*Rev-erbα*^−/−^ mice. The transcriptomic profiles resolved endothelial^45^, erythroid^46^, macrophage/foam cell^3^, smooth muscle cell (SMC), fibroblast, and osteochondroblastic populations (see Table 3 for cell annotation) (Figure 4A, Supplementary Figure 3A-C). Foam cells express *Plin2, Abcg1*, and *Cd68* and, in some cases, *Ptprc*, consistent with contributions from both residual macrophages and SMCs undergoing phenotypic modulation^3^ (Figure 4A, Supplementary Figure 3A-C). Acta2^+^ clusters encompass contractile, proliferative, migratory, and osteochondroblastic SMC states ^13,47^. Contractile SMCs are defined by *Cnn1* and *Myh11*^*48,49*^, whereas loss of these markers accompanies acquisition of proliferative (*Ccnd1, Tff2*)^50,51^ or migratory (*Phactr1*)^11^ phenotypes (Figure 4A, Supplementary figure 3A-C). Osteochondroblastic populations express *Sox9*^*47,52*^ and *Postn*^*53*^ and include pericyte-derived osteoblast-like cells (*Wif1*)^54^, chondroblast-like cells expressing *Spp1, Acan, Tnfsr11b*, and *Col2a1*^*53,55*^, and osteoblast/pre-osteocyte populations marked by *Anxa8, Comp*, and *Cxcl14*^*56*^ (Figure 4A, Supplementary Figure 3A-C). Fibroblast-derived osteogenic cells are identified by co-expression of fibroblast markers (*F3, ApoD, C7*)^53,57,58^, osteoblast-associated genes (*Gdf1*0, *Smoc2*)^59,60^ and chondromyofibroblast-derived markers (*Ptx3, Tmem100, Has1*)^61–63^ (Figure 4A, Supplementary Figure 3A-C).

**Figure 4:**
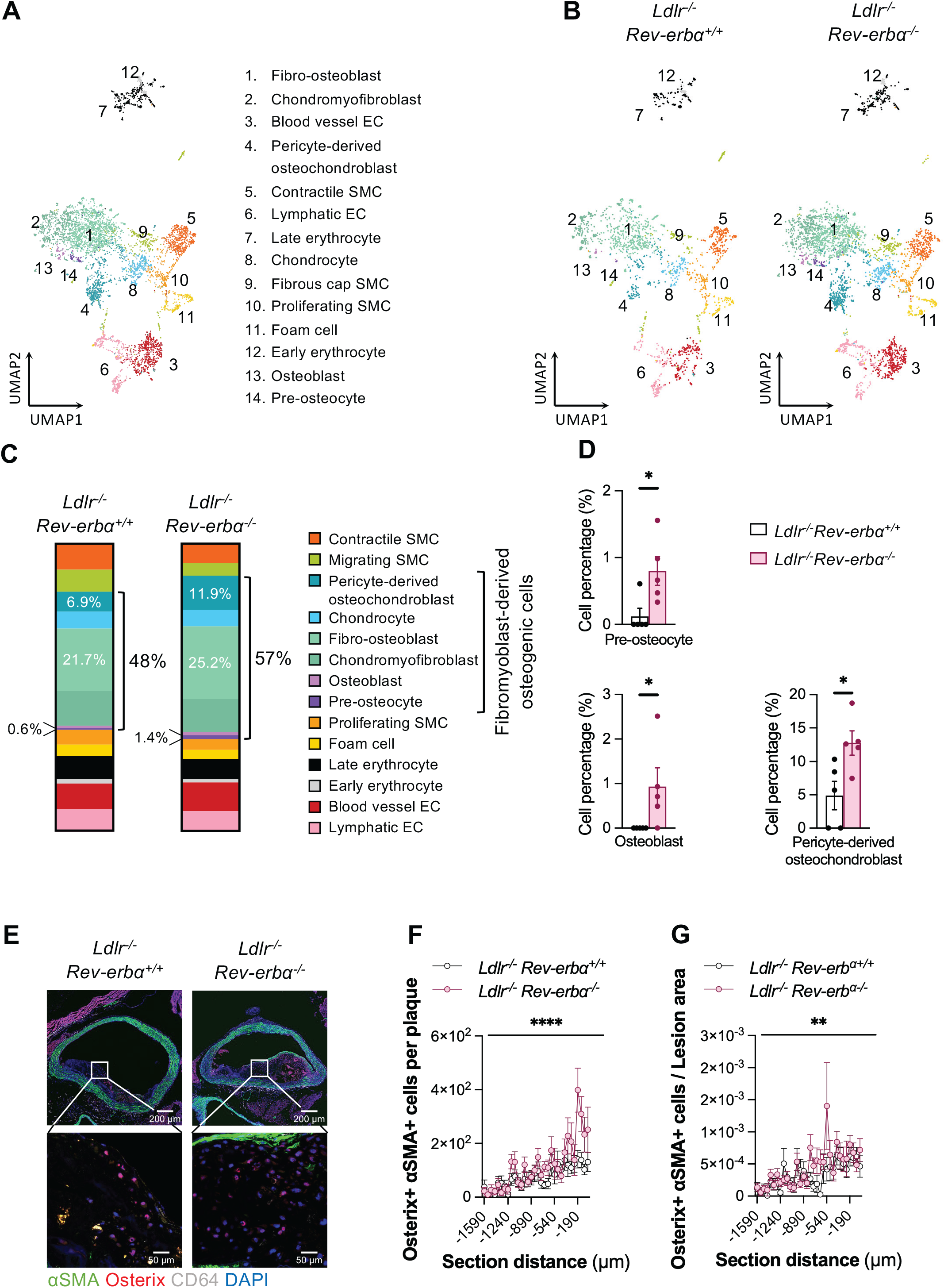
Rev-erbα deficiency promotes osteogenic smooth muscle cell phenotypic transition in atherosclerotic lesions. **A-C**. Single-cell RNA sequencing (scRNA-seq) was performed on cells isolated from lesion areas in the aortas of 18-month-old *Ldlr*^−/−^*Rev-erbα*^*+/+*^ and *Ldlr*^−/−^*Rev-erbα*^−/−^ mice (n=5 per genotype). Uniform Manifold Approximation and Projection (UMAP) plots visualization of CD45-cells, color coded for the identified cell phenotypes in all mice (A) and per genotype (B). Bar graphs showing the relative fraction (%) of each cell type in atherosclerotic plaque from 18-month-old *Ldlr*^−/−^*Rev-erbα*^*+/+*^ and *Ldlr*^−/−^*Rev-erbα*^−/−^ mice (C). **D**. Percentage of different cell types in atherosclerotic plaque from 18-month-old *Ldlr*^−/−^*Rev-erbα*^*+/+*^ and *Ldlr*^−/−^*Rev-erbα*^−/−^ mice (n=5 per genotype) assessed by scRNA sequencing. Statistical analysis was performed using Mann-Whitney test *p < 0.05. **E**. Representative staining of osterix (red), α-SMA (SMC, green), CD64 (macrophages, white) and DAPI (Nuclei, blue) in ascending aortic sections from 18-month-old *Ldlr*^−/−^*Rev-erbα*^*+/+*^ (n=9) and *Ldlr*^−/−^*Rev-erbα*^−/−^ (n=10) mice by confocal microscopy (upper panel: Objectif 10x/air, lower panel: Objectif 40x/1.3 oil). Contrast and brightness adjustments were applied equally to each condition using ImageJ2 software (2.16.0/1.5t). **F-G**. Quantification of osterix-positive smooth muscle cells (F) and normalized to lesion area (G) as a function of location relative to valves. Data are presented as means ± SEM (n= 5-9 per genotype). Two-way ANOVA test ** *p*< 0.01 **** *p*< 0.0001.

Loss of Rev-erbα increases the proportion of osteogenic cells, particularly osteoblasts, pre-osteocytes, and pericyte-derived osteochondroblasts (Figure 4B-D, Supplementary Figure 4). Notably, osteoblasts and pre-osteocytes display a 5-fold increase in *Ldlr*^−/−^*Rev-erbα*^−/−^ mice but represent minor cell types essentially undetectable in *Ldlr*^−/−^*Rev-erbα*^*+/+*^ control mice at the sampling depth. Consistent with these findings, Sp7 (osterix) protein expression is increased in α-SMA^+^ cells of the ascending aorta in *Ldlr*^−/−^*Rev-erbα*^−/−^ mice (Figure 4E-G). Thus, advanced atherosclerotic plaques harbor CD45^−^ cells derived from SMCs and fibroblasts that acquire osteoblast-like phenotypes, and Rev-erbα restrains this pathological osteogenic transition.

### Rev-erbα controls osteoblast differentiation of SMC *in vitro*

To assess whether Rev-erbα plays a functional role in the transdifferentiation of SMC into osteoblast-like cells, we incubated primary SMCs isolated from the aortas of *Rev-erbα*^*+/+*^ and *Rev-erbα*^−/−^ littermate mice in a phosphate-containing osteogenic medium. Calcium deposition is significantly higher in *Rev-erbα*^−/−^ compared to *Rev-erbα*^*+/+*^ SMCs (Figure 5A-B), along with higher activity and expression of Tissue - Nonspecific Alkaline Phosphatase (TNAP) (Figure 5C-D). In line, the expression of osteoblastic markers such as osteocalcin (*OCN*), osteopontin (*Spp1*) and osterix (*Sp7*) are increased in *Rev-erbα*^−/−^ compared to *Rev-erbα*^*+/+*^ SMCs (Figure 5E-G), whereas the anti-calcifying marker Matrix Gla Protein (MGP), a calcium trap, is lower in *Rev-erbα*^−/−^ primary aortic SMCs (Figure 5H). Conversely, PO_4_ ^3-^-challenged primary aortic SMCs treated with hemin, the natural Rev-erbα/β ligand, displayed decreased calcium deposition, osteocalcin and osteopontin expression (Figure 5I-L), whereas MGP expression slightly increased upon hemin treatment (Supplementary Figure 5A). Altogether, our data demonstrate that Rev-erbα inhibits the transcriptional reprogramming of SMCs into osteoblast-like cells.

**Figure 5:**
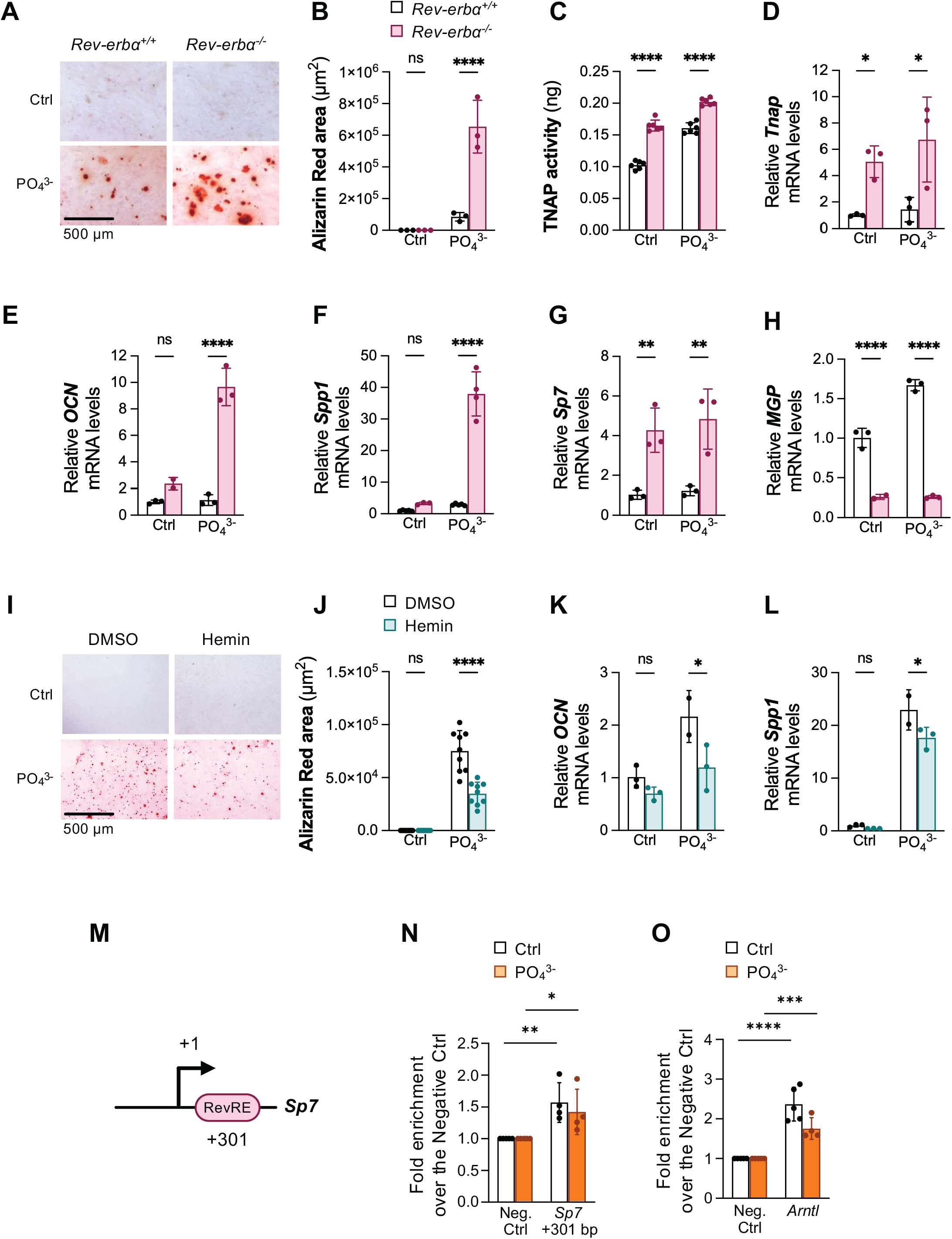
Effect of Rev-erbα deletion and pharmacological activation of Rev-erb-α on osteoblastic differentiation of primary aortic SMCs. **A-B**. Representative staining (A) and quantification (B) of calcium deposits after alizarin red staining in SMC isolated from Rev-erbα^+/+^ and Rev-erbα^−/−^ mice cultured with or without osteogenic differentiation medium (Na_2_PO_4_.2H_2_O, 3 mM) for 15 days (12 fields per well, n=3 per condition). Data are represented as mean ± SD. Contrast and brightness adjustments were applied equally to each condition using ImageJ2 software (2.16.0/1.5t). Statistical analysis was performed using a two-way ANOVA test followed by a Šídák post-hoc test. **** p<0.0001 **C**. Tissue non-specific alkaline phosphatase (TNAP) activity assay (n=6 per condition). Data are means ± SD. Statistical analysis was performed using a two-way ANOVA test followed by a Šídák post-hoc test. **** p<0.0001. **D-G**. mRNA expression levels of the pro-calcifying osteoblastic markers Tnap (tissue nonspecific alkaline phosphatase), Bglap (osteocalcin), Spp1 (osteopontin) and Sp7 (osterix). **H**. mRNA expression levels of the anti-calcifying osteoblastic markers Mgp (Matrice Gla Protein). Data are means ± SD (n=3 per condition). Statistical analysis was performed using a two-Way ANOVA test followed by a Šídák post-hoc test. * p<0.05, ** p<0.01, **** p<0,0001. **I-J**. Representative staining (I) and quantification (J) of calcium deposits after alizarin red staining in SMC isolated from Rev-erbα^+/+^ mice treated or not with hemin (6µM) and cultured or not with osteogenic differentiation medium (Na_2_PO_4_, 2H_2_O 3 mM) for 15 days (12 fields per well, n=3 per condition). Contrast and brightness adjustments were applied equally to each conditions using ImageJ2 software (2.16.0/1.5t). Data are represented as mean ± SD. Statistical analysis was performed using a two-way ANOVA test followed by a Šídák post-hoc test. **** p<0.0001. **K-L**. mRNA expression levels of the osteoblastic markers Bglap (osteocalcin) and Spp1 (osteopontin). Data are mean ± SD (n=3 per condition). Statistical analysis was performed using a two-way ANOVA test followed by a Šídák post-hoc test. * p<0.05. **M**. Diagram illustrating putative Rev-erb-α response elements (RevRE) in the Sp7 promoter, obtained using MatInspector software. **N**-**O**. Chromatin immunoprecipitation (ChIP) with an antibody directed against Rev-erb-α on the promoter of Sp7 (N) and Arntl (Bmal1) (O) were performed in MOVAS synchronized and cultured or not with osteogenic differentiation medium (Na_2_PO_4_, 2H_2_O 3 mM) (n=4 per condition). Data represent means ± SD. Statistical analysis was performed using a two-way ANOVA test followed by a Šídák post-hoc test. * p<0.05, ** p<0.01, *** p<0.001, **** p<0.0001.

We thus assessed whether Rev-erbα directly binds to the gene promoters of the regulated osteogenic transcription factors to inhibit their expression. *In silico* analysis identified several putative Rev-erb Response Elements (RevRE) in the *Sp7* (Figure 5M) and *Spp1* promoters (Supplementary Figure 5B). Chromatin Immunoprecipitation assays demonstrate that Rev-erbα binds to a RevRE located +301 pb in the 5’UTR of the *Sp7* gene (Figure 5N), as well as to RevREs located at −6700 pb, −6520 bp, −3070 bp, −2120 bp and −560 bp in the *Spp1* promoter (Supplementary Figure 5C-G). The RevRE site located in the *Arntl* (Bmal1) gene was used as a positive control (Figure 5O, Supplementary Figure 5H). In conclusion, Rev-erbα prevents the differentiation of SMCs into osteoblasts by inhibiting the transcription of key osteoblastic transcription factors through direct binding to their promoters.

### Rev-erbα represses the NLRP3 inflammasome pathway and IL-1β-induced calcification

Since the production of pro-inflammatory cytokines, including IL-1β, accelerates the dedifferentiation of SMC and since Rev-erbα inhibits the NLRP3-mediated IL-1β secretion and maturation in macrophages^24^, we assessed whether this mechanism is also operative in the context of atherosclerosis. In human endarterectomies from the OPAL cohort, *NLRP3* expression is elevated in lesions compared to healthy areas (Supplementary Figure 6A), concomitant with lower *REV-ERBα* expression (Figure 1E). In aortas from 18-month-old *Ldlr*^−/−^*Rev-erbα*^−/−^ mice, the expression of *Nlrp3* and the pro-inflammatory cytokines *IL-1β* and *IL-18* is up-regulated compared to aortas from *Ldlr*^−/−^*Rev-erbα*^*+/+*^ littermate controls (Supplementary Figure 6B). Furthermore, NLRP3 protein levels are higher in macrophages, but not in SMCs, located within the lesions of *Ldlr*^−/−^*Rev-erbα*^−/−^ mice compared to their littermate controls (Supplementary Figure 6C-D). To investigate whether Rev-erbα also inhibits the IL-1β-induced SMC dedifferentiation into osteoblast-like cells, we incubated primary aortic SMCs from *Rev-erbα*^*+/+*^ and *Rev-erbα*^−/−^ mice in a PO_4_^3-^-containing calcifying medium and then challenged them with IL-1β. IL-1β strongly exacerbated the effect of *Rev-erbα* deficiency on calcium deposition in primary aortic SMCs cultured under osteogenic conditions (Supplementary Figure 6E-F). Collectively, these findings indicate that Rev-erbα regulates NLRP3 signaling in human and mouse atherosclerotic plaques and modulates SMC sensitivity to IL-1β-induced osteoblastic differentiation *in vitro*.

**Figure 6:**
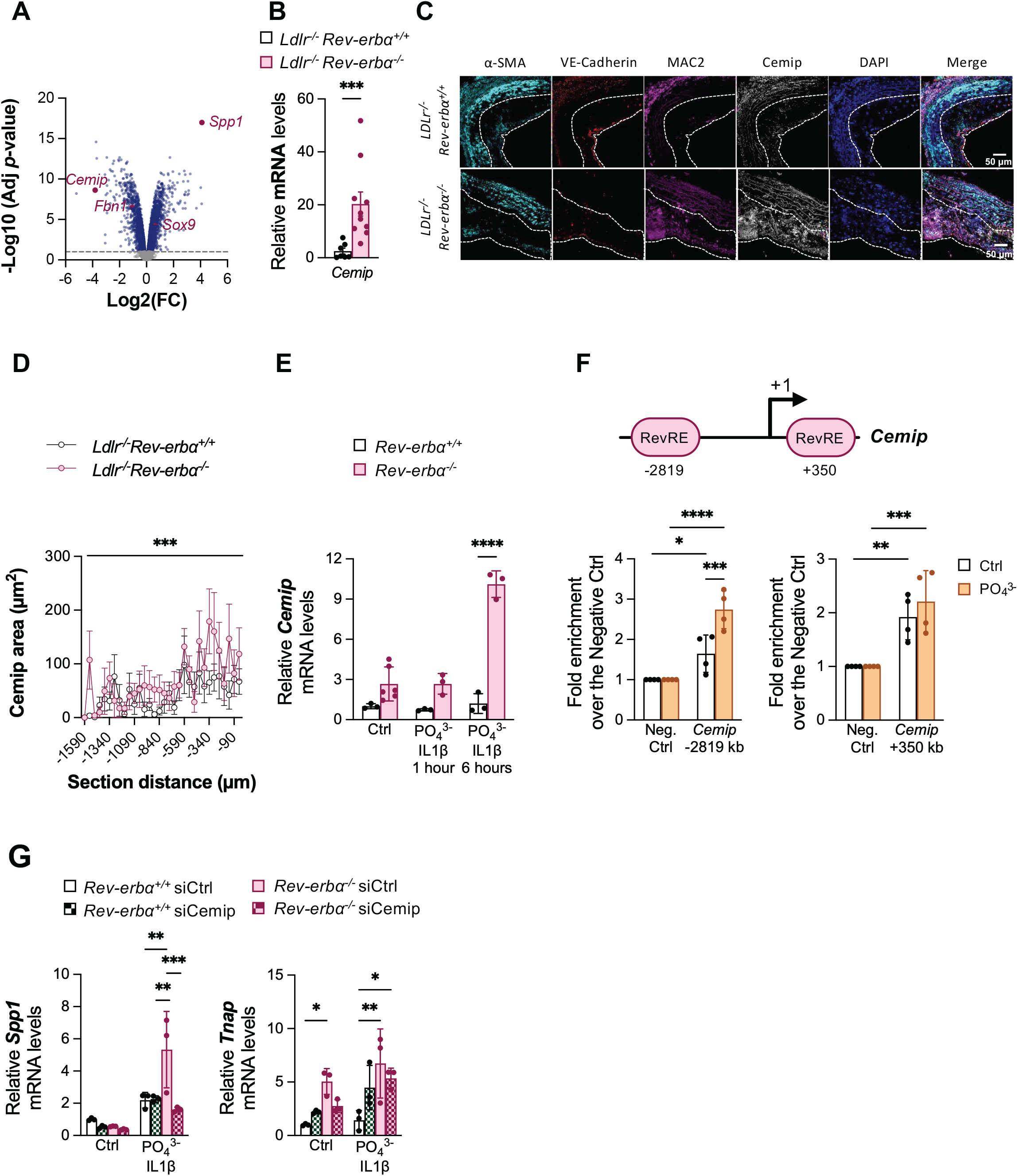
Impact of Rev-erbα deletion on Cemip expression in atherosclerotic plaques and osteoblastic differentiation. **A**. Volcano-plot representation of a representative metabolic pathway analysis of calcification pathway enrichment performed on isolated SMCs from Rev-erbα^+/+^ and Rev-erbα^−/−^ mice cultured with or without osteogenic differentiation medium (Na_2_ PO_4_ ·2H_2_ O, 3 mM) for 15 days (n= 3-5 per condition as indicated). **B**. Cemip mRNA expression levels in whole aortas isolated from Ldlr^−/−^Rev-erbα^+/+^ and Ldlr^−/−^Rev-erbα^−/−^ mice. Data are shown as means ± SEM (n= 7-10 per genotype). Statistical analysis was performed using a Mann-Whitney test. *** p<0.001. **C-D**. Representative Cemip immunofluorescence staining (grey) (C) and quantification of the total number of Cemip positive cells (D) in different atherosclerotic plaque cells (SMC: αSMA cyan; endothelial cells: Ve-cadherin red; macrophages: MAC2 purple) based on their location relative to valves in histological sections of ascending aortas from 18-month-old Ldlr^−/−^Rev-erbα^+/+^ (n=8) and Ldlr^−/−^ Rev-erbα^−/−^ (n=9) mice. Data are represented as means ± SEM. Statistical analysis was performed using a two-way ANOVA test followed by a Šídák post-hoc test. ** p<0.01, *** p<0.001. **E**. Cemip expression level in SMCs isolated from Rev-erbα^+/+^ and Rev-erbα^−/−^ after acute treatment with IL-1β mice (n=3 per condition). Data are means ± SD. Statistical analysis was performed using a two-way ANOVA test followed by a Šídák post-hoc test. **** p<0.0001. **F**. Representation of putative Rev-erb-α response elements (RevRE) in the Cemip promoter obtained using the EPD interface and ChIP analysis of Rev-erbα occupancy of the Cemip promoter in MOVAS cells, synchronized and cultured with or without osteogenic differentiation medium (Na_2_ PO_4_ ·2H_2_ O, 3 mM) (n=4 per condition). Data represent means ± SD. Statistical analysis was performed using a two-way ANOVA test followed by a Šídák post-hoc test. * p<0.05, ** p<0.01, *** p<0.001, **** p<0.0001. **G**. Spp1 and Tnap mRNA expression levels in SMC isolated from Rev-erbα^+/+^ and Rev-erbα^−/−^ mice, with or without Cemip expression inhibition by siRNA, and cultured with or without osteogenic differentiation medium (Na_2_ PO_4_ ·2H_2_ O, 3 mM) ± IL-1β (0.1 ng/mL) for 48h (n=3 per condition). Data are shown as mean ± SD. Statistical analysis was performed using a two-way ANOVA test followed by a Šídák post-hoc test. ** p<0.01, *** p<0.001.

### Rev-erbα regulates the expression of the hyaluronidase Cemip during vascular calcification

To further characterize genes whose expression is modulated by Rev-erbα during SMC osteogenic differentiation, we conducted a transcriptomic profiling of primary aortic SMC from *Rev-erbα*^*+/+*^ and *Rev-erbα*^−/−^ mice cultured under either standard or osteogenic conditions. PCA analysis and identification of differentially expressed genes (DEGs) confirmed distinct transcriptomic signatures across the 4 groups, revealing both genotype and treatment effects (Supplementary Figure 7A-B). Interestingly, osteogenic stimulation affects common and distinct sets of upregulated genes depending on the genotype (Supplementary Figure 7C). We then sought to identify differentially expressed genes and enriched pathways whose regulation during osteoblastic differentiation is affected by the absence of Rev-erbα. Using a statistical interaction analysis approach^33^, we found that the expression of genes directly and indirectly involved in osteoblast differentiation, including *Spp1, Sox9, Fbn1*, is altered by the deletion of Rev-erbα in phosphate-challenged SMC (Figure 6A). Consistently, fGSEA revealed these genes were particularly enriched in pathways controlling osteogenesis such as osteoblast differentiation and development, cartilage development, bone mineralization and regulation of the Wnt and the Notch signaling pathway, which both drive osteoblastic differentiation and promote SMC calcification^64,65^ (Supplementary Figure 7D). Intriguingly, the extracellular matrix reorganization process was also altered by the deletion of Rev-erbα in phosphate-challenged SMC (Supplementary Figure 7D). Accordingly, we identified the cell migration inducing protein (Cemip) to be dysregulated between *Rev-erbα*^*+/+*^ and *Rev-erbα*^−/−^ primary aortic SMC undergoing osteoblastic differentiation (Figure 6A). We tnen confirmed that Cemip expression is increased at the mRNA and the protein levels in ascending aortic atherosclerotic lesions from *Ldlr*^−/−^*Rev-erbα*^−/−^ mice (Figure 6B-D), which is in line with the enrichment in genes of the hyaluronan degradation pathways observed in 18-month-old *LDLr*^−/−^*Rev-erbα*^−/−^ aortas (Figure 3E). *In vitro* experiments confirmed that *Rev-erbα* deficiency results in higher *Cemip* expression within SMCs under inflammatory and osteogenic conditions (Figure 6E). Finally, ChIP experiments demonstrate that Rev-erbα is recruited to two RevREs located in the *Cemip* promoter approximately 2.8 kb and 0.3 kb upstream from the transcription start site (Figure 6F), while Rev-erbα binds to the *Arntl* promoter used as a positive control (Supplementary Figure 7E). Interestingly, the binding of Rev-erbα to the −2.8 kb RevRE in the *Cemip* promoter is further increased in osteogenic conditions (Figure 6F). Finally, while *Spp1* and *Tnap* gene expression are increased in *Rev-erbα*^−/−^ primary aortic SMC cultured under inflammatory and osteogenic conditions compared to *Rev-erb*^*+/+*^ SMC, *Cemip* knock-down abolishes the effect of Rev-erbα deficiency on these SMC transdifferentiation markers, thus demonstrating that Rev-erbα effect on osteogenic differentiation depends, at least partially, on the regulation of *Cemip* expression (Figure 6G, Supplementary Figure 7F). Altogether, these results identify an instrumental role of Rev-erbα in phosphate-triggered SMC-to-osteoblast transition and identify the hyaluronan degradation pathway, and particularly Cemip, as a Rev-erbα target to combat osteoblast differentiation and vascular calcification.

## Discussion

Combining studies on human samples, preclinical models and primary cells, we identify the nuclear receptor Rev-erbα as a central regulator of vascular calcification (Supplementary Figure 8). Single-cell profiling of whole aortas as well as *in vitro* experiments in primary aortic SMCs revealed an important role of Rev-erbα in SMC osteoblastic transdifferentiation. Interestingly, Pan *et al* observed that a decrease in Rev-erbα (Nr1d1) is a transcriptional signature of SMC transition toward SMC-derived intermediate cells in mouse atherosclerotic plaques, in line with our finding of the important role of Rev-erbα expression in maintaining SMC identity during atherogenesis^13^. Accordingly, *Rev-erbα* deficiency modulates the SMC response to inflammatory stimuli by directly shaping a transcriptomic reprogramming towards transdifferentiation into osteoblast-like cells. Moreover, whole aorta and *in vitro* cell transcriptomic analysis demonstrated that Rev-erbα regulates genes involved in hyaluronic acid degradation and matrix remodeling pathways, thereby counteracting plaque progression and vascular calcification. Given that Rev-erbα can be activated by synthetic ligands, the development of pharmacological modulators may represent a potential strategy to prevent atherosclerosis progression and vascular calcification.

Interestingly, we now report that *REV-ERBα* expression is particularly lower in calcified areas compared to adjacent non-calcified plaque regions. This extends a previous study in mice showing that *Rev-erbα* deficiency increases intraplaque hemorrhage and ECM remodeling in *ApoE*^−/−^ mice, thereby increasing the risk of plaque instability and rupture^66^. Calcification is an active process resulting from an imbalance between osteoclastic-mediated resorption and osteoblastic-induced mineralization differentiation. Interestingly, Rev-erbα was shown to control the balance in bone metabolism, primarily through inhibition of the p38 MAPK pathway^67^. In addition, Rev-erbα was suggested to suppress the osteoblastic differentiation by preventing RUNX2 nuclear translocation and the subsequent induction of its target genes^67^. Consistently, we observed higher expression of *Runx2* in *Ldlr*^−/−^*Rev-erbα*^−/−^ aortas compared with controls. Our data further indicate that Rev-erbα coordinately regulates several key osteogenic genes, including *Spp1* and *Sp7*. Osteopontin (*Spp1*) is of particular interest because it promotes plaque progression^68^ by increasing lipid streak formation^69,70^ and by regulating SMC proliferation and migration^71,72^. Furthermore, osteopontin was shown to be elevated in calcified aortas and to control vascular calcification^73,74^. Accordingly, we identify osteopontin as a novel Rev-erbα target gene in atherosclerotic plaques, suggesting that Rev-erbα counteracts osteopontin pro-calcifying and pro-atherogenic actions. Similarly, Rev-erbα represses osterix (*Sp7*) expression, a key transcription factor that acts as a molecular switch in osteoblastic differentiation. Osterix-null mice exhibit impaired bone formation due to defective bone matrix deposition^75,76^, emphasizing the importance to tightly regulate osterix expression. Beyond bone metabolism, osterix plays a role in mineralization processes in several contexts, including periodontitis, a chronic inflammatory disease affecting tooth-supporting tissues and associated with an increased risk of atherosclerosis^77^. Consistent with our observations, elevated *Rev-erbα* expression has been reported in senescent cementoblasts, where it suppresses cementoblast proliferation and impairs mineralization in an osterix-dependent manner^77^. We demonstrate that Rev-erbα directly binds the promoter of *Sp7* and *Spp1* to repress their transcription, offering a mechanistic basis for our observations. Consequently, the absence of Rev-erbα leads to a de-repression of these genes, as well as TNAP, *Sox9* and *Bglap*, allowing SMCs to transdifferentiate into osteoblast-like cells and the formation of calcium deposits. Besides, our study further reveals that Rev-erbα exerts a broader impact on vascular SMC plasticity, shaping their identity and fate during atherosclerosis progression. More specifically, Rev-erbα deficiency drives SMC and pericyte differentiation toward an osteoblast phenotype. Conversely, pharmacological activation of Rev-erbα with hemin, its natural ligand, reduces the osteoblastic differentiation of SMCs, suggesting its potential as a therapeutic target for limiting vascular calcification.

Interestingly, the increased calcium deposition observed in *Rev-erbα* deficient mice is localized in the intima, which is particularly relevant given that intimal calcification is a key driver of plaque instability and rupture. On the other hand, the clock component Bmal1, a Rev-erbα target gene, promotes medial, but not intimal, calcification in streptozotocin-induced diabetic mice by inducing *Runx2* expression in SMC^78^. Although we did not observe an effect of Rev-erbα on medial calcification in our *Ldlr*^−/−^ model challenged with a warfarin-containing diet, known to trigger medial calcification^79^ (data not shown), we confirm here a role of the circadian machinery on vascular calcification, but with potentially distinct roles of different clock components in the intima *vs* the media. Transcriptomic analysis of whole aortas from *Ldlr*^−/−^ mice revealed that *Rev-erbα* deficiency is associated with an enrichment of matrix remodeling pathways, particularly those related to hyaluronan degradation. Consistently, transcriptional analysis of primary aortic SMCs confirmed that *Rev-erbα* deficiency activates pathways involved in matrix degradation under osteogenic conditions. This approach also identified the hyaluronidase Cemip as a key mediator of Rev-erbα’s action in driving the osteoblastic differentiation of SMCs. Supporting these findings, inhibition of hyaluronic acid synthesis has been shown to aggravate atherosclerotic lesions^80^, trigger osteogenic marker expression and promote SMC calcification^21^. Conversely, high-molecular-weight hyaluronic acid was reported to actively suppress SMC osteogenic differentiation and calcification^21^. Our results reveal that *Rev-erbα* deficiency upregulates Cemip both *in vivo* and *in vitro* and that knock-down of Cemip in SMC impairs the effect of Rev-erbα deletion on SMC-derived osteoblast differentiation, establishing Cemip as a key downstream effector of Rev-erbα in vascular calcification and matrix remodeling. Cemip has already been shown to contribute to atherosclerosis by promoting SMC proliferation and migration^20^ and by degrading hyaluronic acid into low-molecular-weight fragments that drive inflammation, angiogenesis and cell migration^81^. However, its role in the phenotypic switch of SMC toward an osteochondrogenic fate was not established. This study suggests that the derepression of *Cemip* upon Rev-erbα deficiency further increases the expression of osteogenic markers/drivers, including Spp1, thereby linking ECM remodeling to osteogenic reprogramming.

In conclusion, Rev-erbα is a key regulator of SMC-induced vascular calcification *in vivo* and *in vitro*. Rev-erbα may therefore represent a new therapeutic target to reduce residual risk in patients at high cardiovascular risk by reducing the inflammatory response, osteoblastic transdifferentiation processes.

## Acknowledgements

We thank the BICeL Facility for access to systems and technical advice. We are grateful to Dr Kevin Ory (UMR 1011) for its helpful insight in the setup of the single cell experiment.

## Supplemental material

Supplemental Materials and Methods

Supplementary Figure S1-S8

Major resources tables

References 27-35

