## Supplemental Figures for "Rev-erbα inhibits vascular calcification by repressing osteogenic reprogramming in atherosclerosis"

Supp. Figure 1

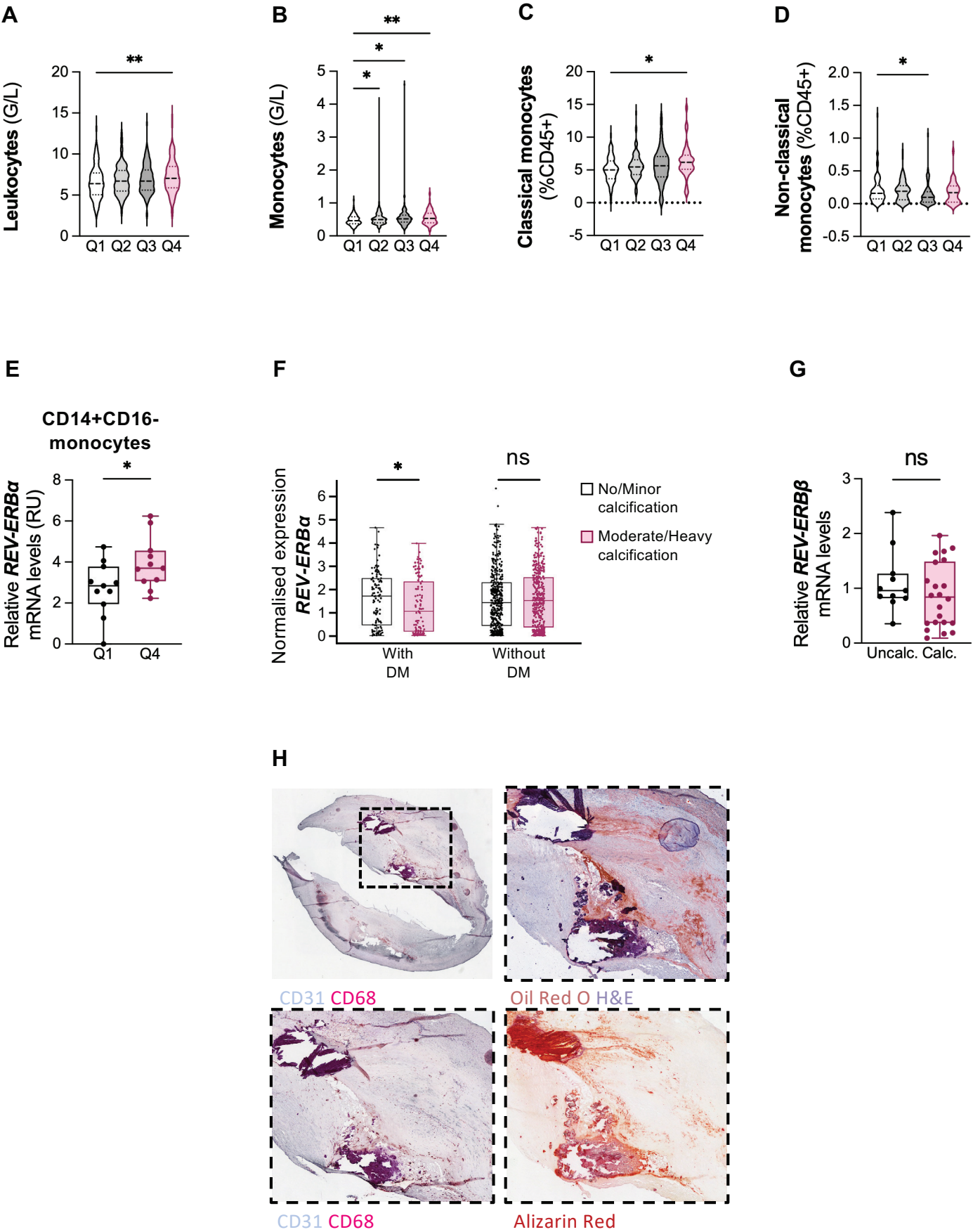

Supp. Figure 2

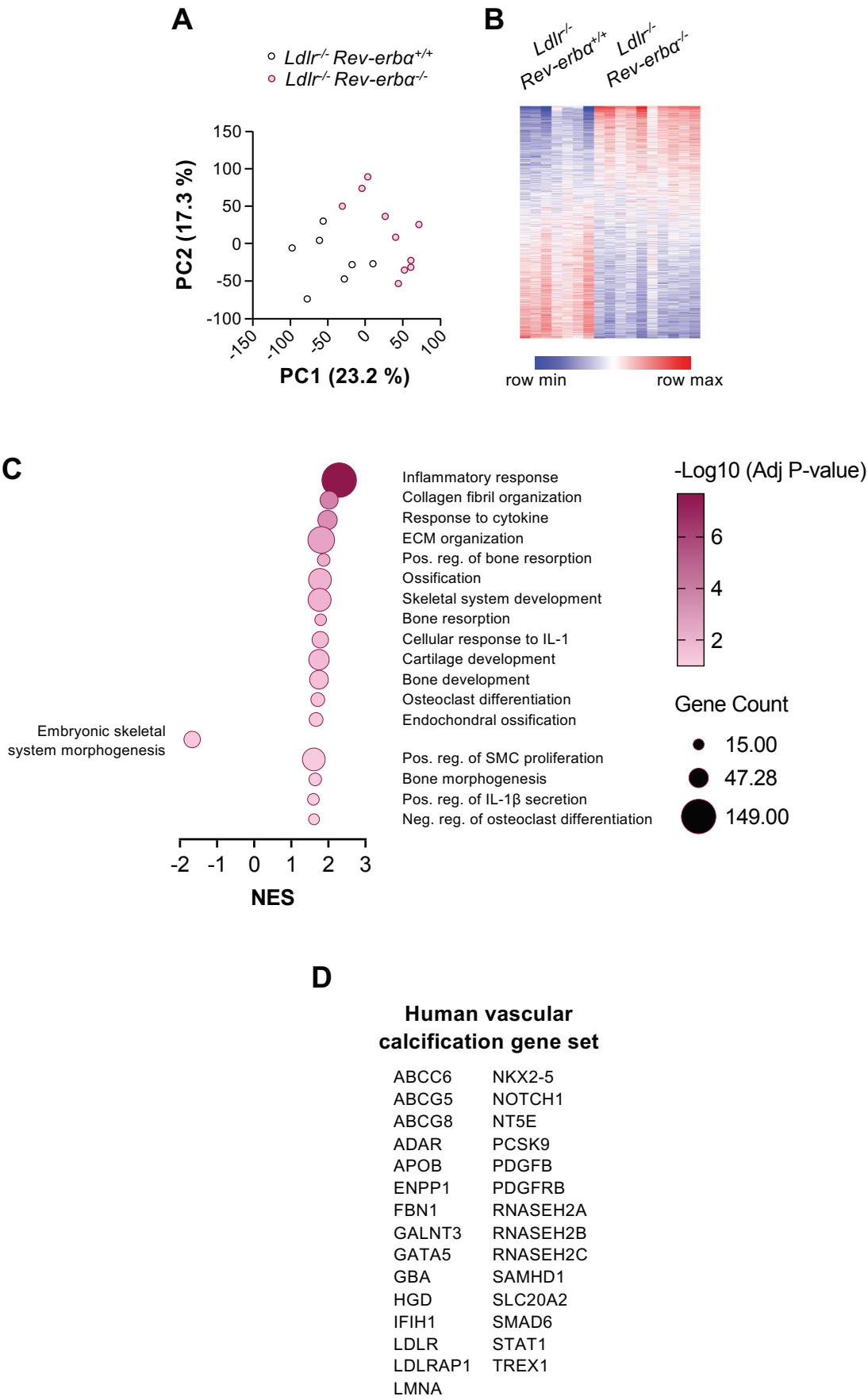

Supp. Figure 3

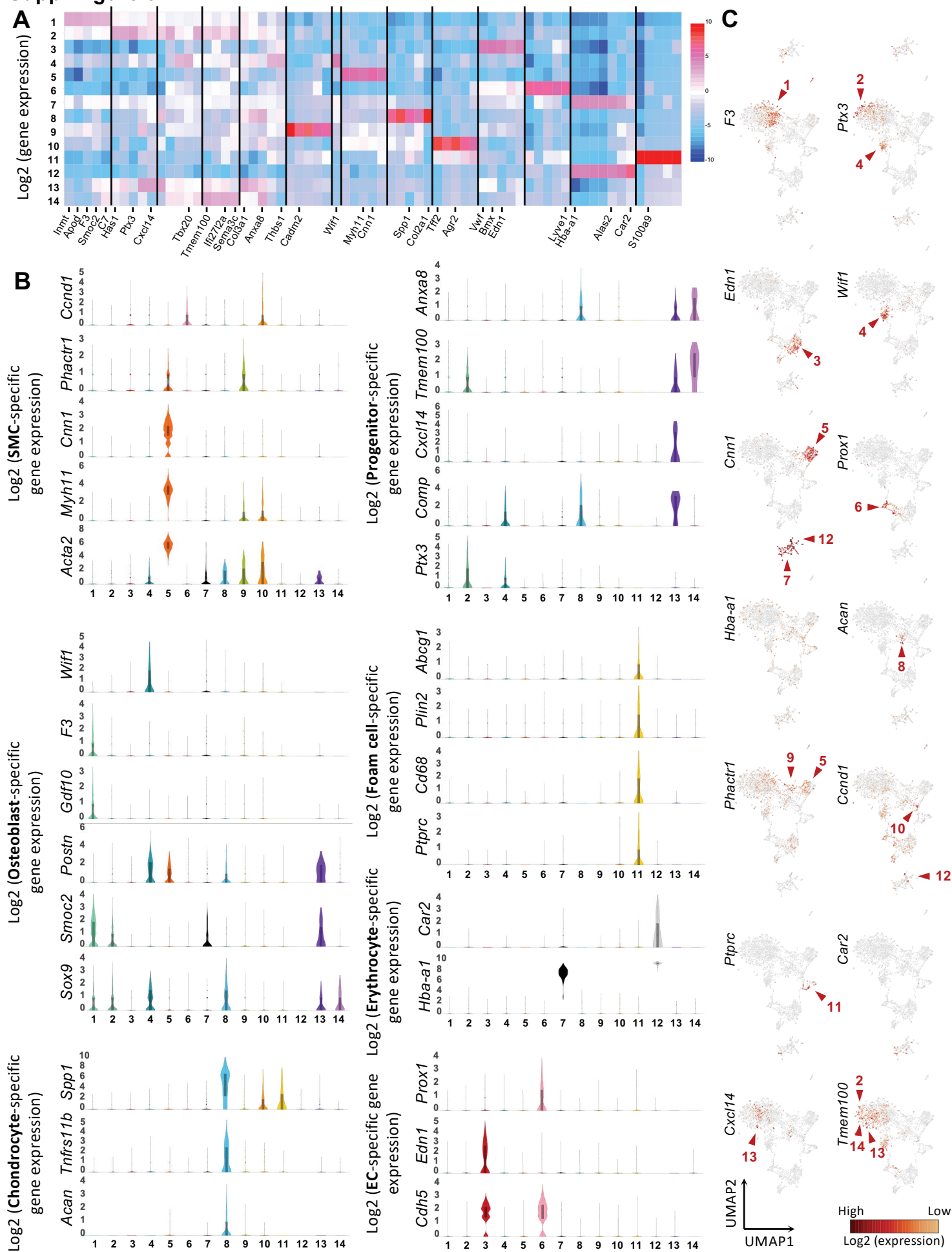

Supp. Figure 4

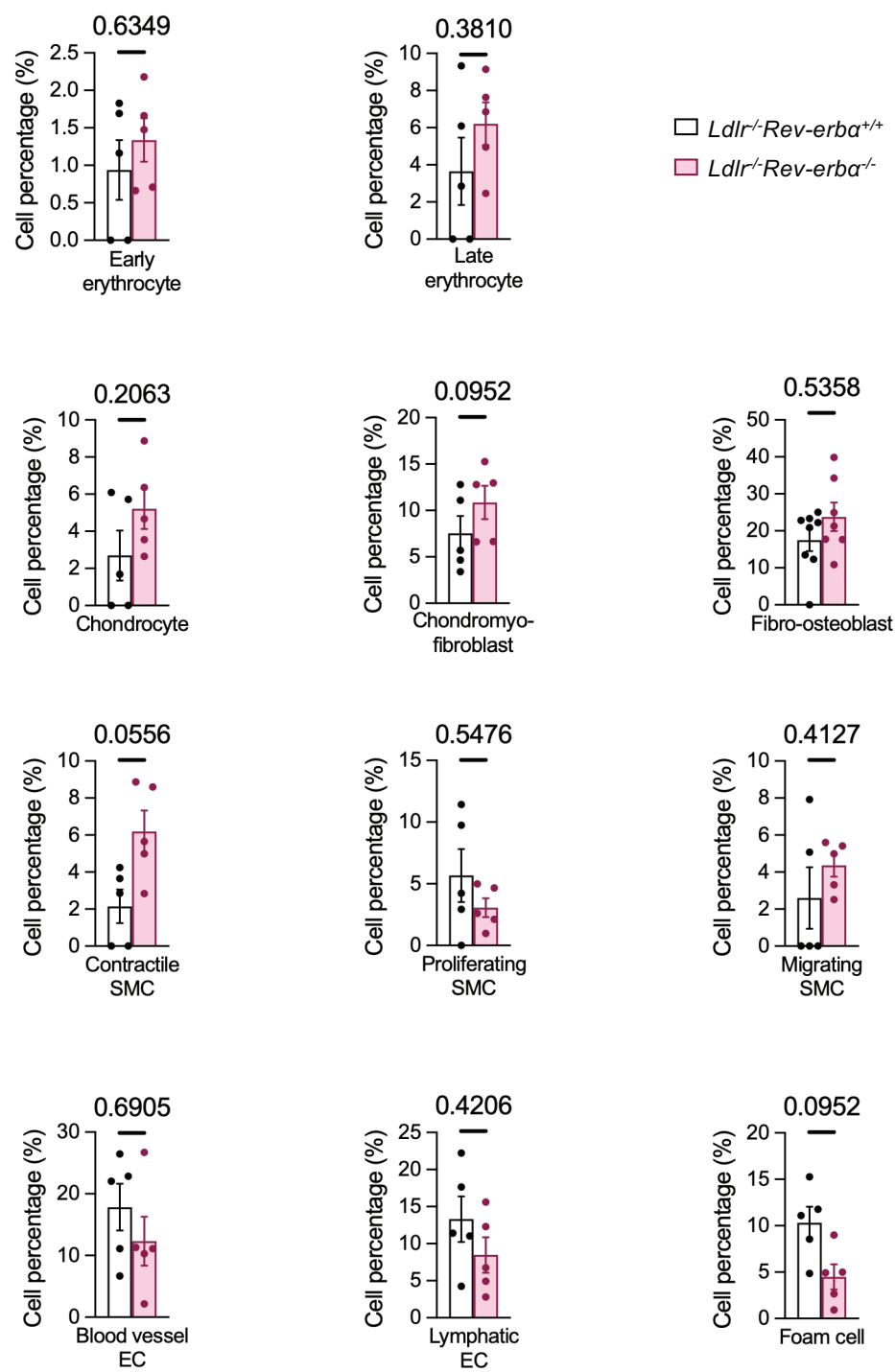

Supp. Figure 5

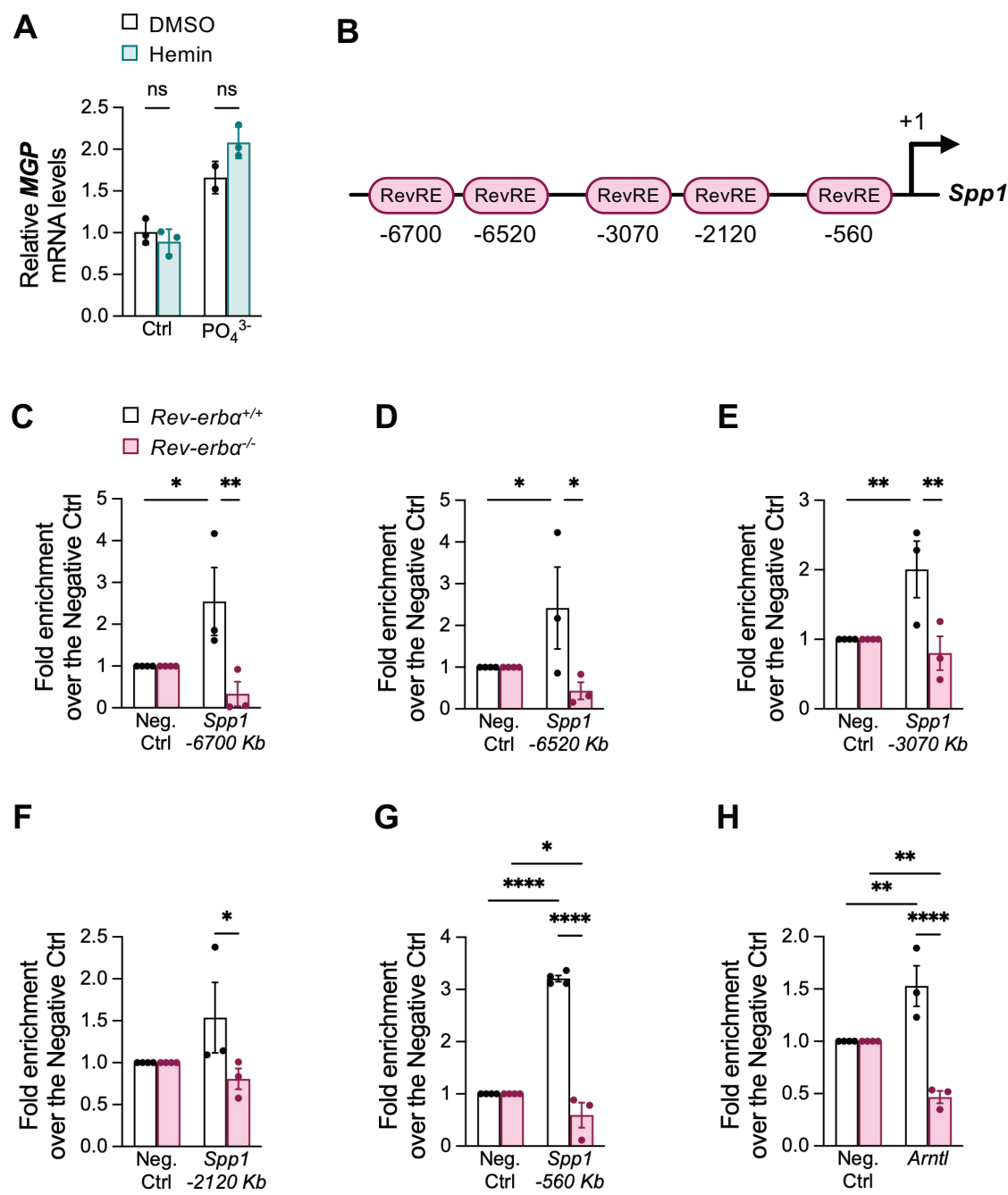

Supp. Figure 6

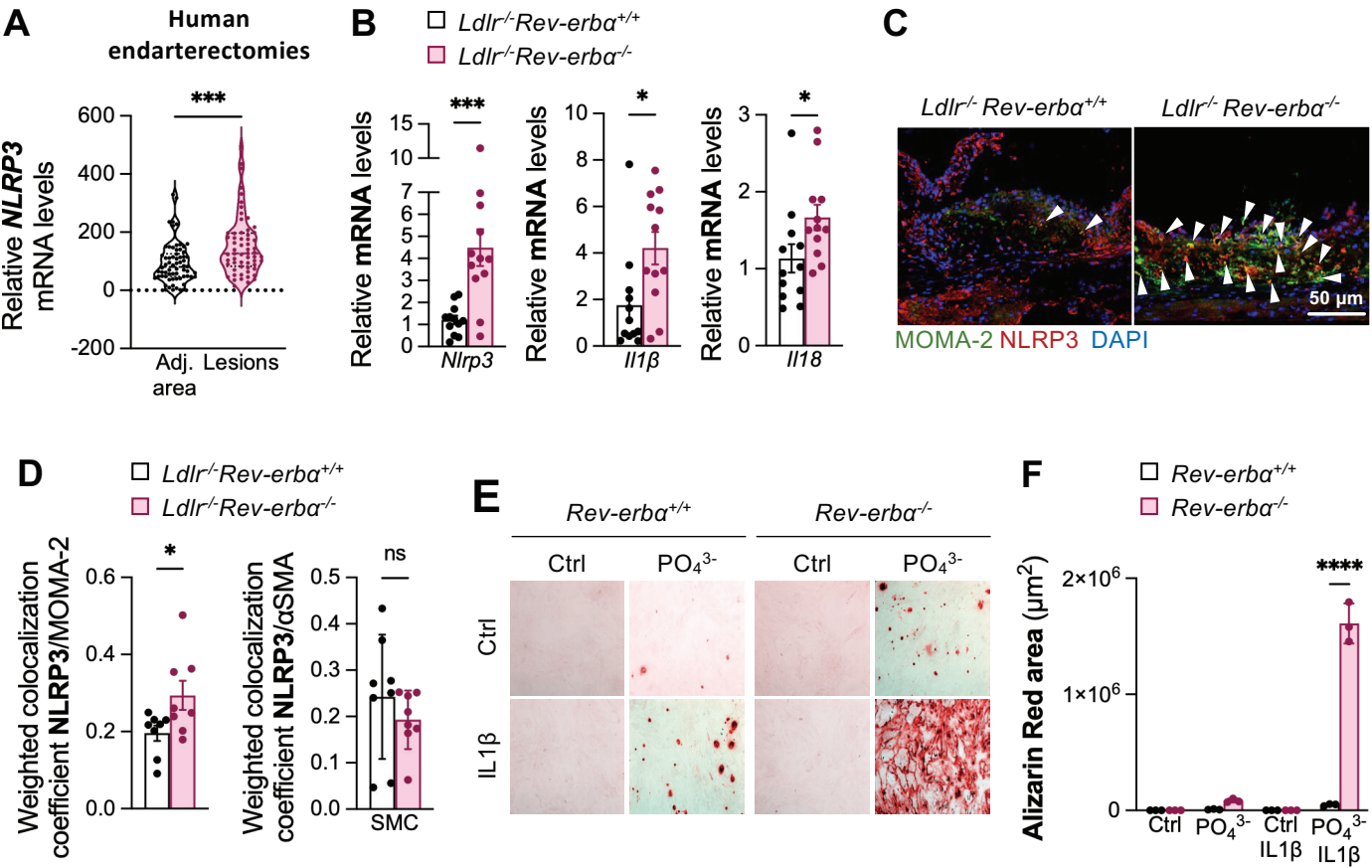

### Supp. Figure 7

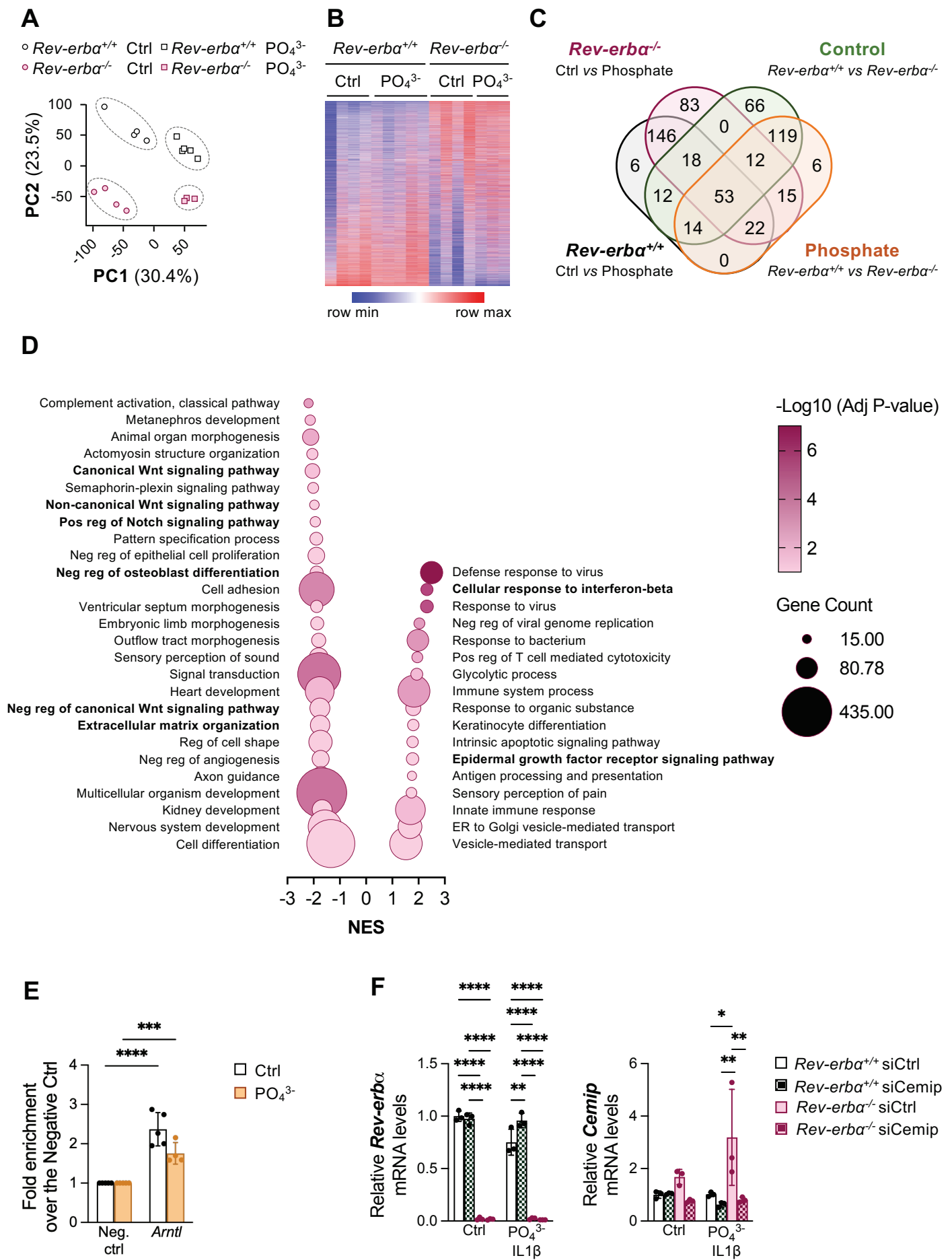

Supp. Figure 8

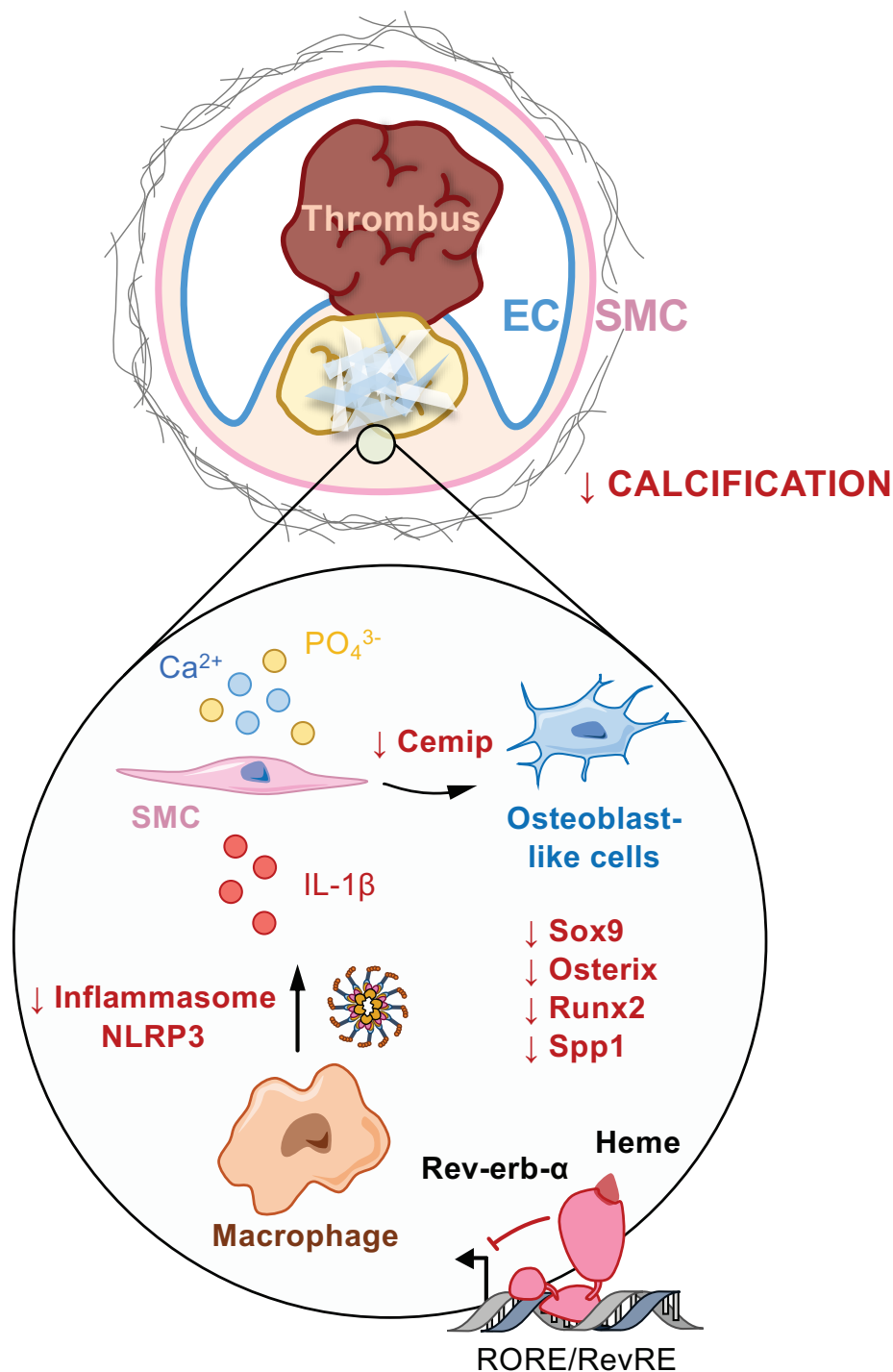
