## Supplemental Material for "Rev-erbα inhibits vascular calcification by repressing osteogenic reprogramming in atherosclerosis"

**Materials and methods**

**Human cohorts**

**ANGIOSAFE cohort**

We analyzed participants from the AngioSafe-2 cohort (ClinicalTrials.gov Identifier: NCT02671864). A total of 668 participants with type 2 diabetes (T2D) were stratified according to their cardiovascular risk using the Coronary Artery Calcium (CAC) score: low CAC (Q1, n=218), moderate CAC (Q2, n=220; Q3, n=128), and high CAC (Q4, n=102). Peripheral blood samples were collected to quantify immune cell populations and circulating anti-inflammatory CD14⁺ monocytes were isolated for transcriptomic profiling. All participants provided informed consent. The study was approved by the relevant institutional review boards and ethics committees, including the CPP Île-de-France Ethics Committee, which authorized the inclusion of all individuals.

**CD14^+^ monocyte sorting and RNA extraction**

20 mL of whole blood was drawn from males with T2D, and PBMCs were isolated using Ficoll Paque^+^. Circulating CD14+ monocytes were immunoselected using anti-human CD14 Milteny MicroBeads. A minimum of 2 million cells were lysed with RLT plus buffer and 3% β-mercaptoethanol for subsequent RNA extraction using the RNeasy Plus Mini extraction kit (Qiagen, Cat. No. 74134), following the user manual instructions. Libraries were prepared using the NEBNext® Ultra™ II Directional RNA Library Prep Kit for Illumina (NEB, E7760L) and paired-end (2x75) strand specific sequencing was performed on Nextseq500 sequencer (Illumina) in two batches at the UVSQ Genomic Platform (Montigny-le-Bretonneux, France). FASTQ files were pre-processed using fastp v0.19.11. Reads were mapped to the GRCh38 primary human genome assembly version 38 with STAR v2.738. Genes with at least 5 raw reads in more than 25% of samples were retained. Reads were assigned to Gencode version 32 features using featureCounts from subread-1.6.4 package39. Differentially expressed genes between CD14+ monocytes from patients in the highest (Q4) and lowest (Q1) coronary artery calcium score quartiles were identified using DESeq2, while accounting for sequencing batch. Gene Ontology over-representation analyses were then performed on significant differentially expressed genes using the Biological Process and Molecular Function collections. Normalization and differential gene expression (DGE) analyses were performed with R package DESeq2 v1.32.0, and Benjamini-Hochberg correction was applied with a threshold for significance set at: adjusted *p-value*s < 0.05.

**OPAL Lille cohort**

Human atherosclerotic plaques were removed from patients eligible for surgical carotid endarterectomy (significant carotid artery stenosis >70%) recruited at the Cardiovascular Surgery Department, Lille University Hospital, France^27^. The study protocol was approved by the local ethics committee (OPAL-Lille study, CPP11-15 n°2010-A01401-38) in accordance with the ethical guidelines of the Declaration of Helsinki. Written informed consent was obtained from all subjects.

Immediately after endarterectomy, representative parts of carotid plaques were embedded in Neg-50 frozen section medium (Microm International, Waldorf, Germany), frozen in liquid nitrogen and stored at −80°C. Patient data were prospectively collected and included cardiovascular medications and risk factors such as sex, age, diabetes mellitus (fasting plasma glucose ≥1.26 g/L and/or use of glucose-lowering medications), dyslipidemia (serum triglyceride levels ≥1.5 g/L or high-density lipoprotein (HDL) cholesterol <0.4 g/L in men or <0.5 g/L in women, and/or current treatment), obesity (body mass index (BMI)) and smoking history (current or former smokers *vs* non-smokers). Additional data included history of myocardial infarction, stroke and peripheral arterial disease (Fontaine claudication stage ≥IIa and/or history of treatment for chronic limb ischemia), as well as left ventricular ejection fraction. Patients were included during outpatient consultation, remote from any surgical procedure. At inclusion, none of the patients presented clinical symptoms of systemic inflammation.

Immunohistochemical analysis was performed on serial 10-μm cryosections after quenching of endogenous peroxidase activity. Vascular SMC (VSMC), endothelial cells and macrophages were identified using antibodies against α-smooth muscle actin (α-SMA), CD31/PECAM-1 (platelet endothelial cell adhesion molecule-1; Novus Biological, Littleton, CO, USA) and CD68 antibodies (Dako), respectively, and N-Histofine Simple Stain (Nichirei Biosciences Inc., Tokyo, Japan). Alizarin red S (Sigma-Aldrich, France) staining was performed to detect calcium deposits. Sections of atherosclerotic plaques were microdissected to isolate areas positive (calcified) or negative (non-calcified) to alizarin red staining. Laser capture microdissection was performed on an ArcturusXT Microdissection Instrument with CapSure Macro LCM Caps (MDS Analytical Technologies, Sunnyvale, CA, USA) and areas were captured from 4 adjacent 8-µm sections and pooled for RNA extraction.

**AtheroPortal**

To assess the expression of *REV-ERBα* according to diabetes mellitus status and plaque calcification, we analyzed bulk RNA-sequencing data from the Athero-Express Biobank Study, an ongoing prospective longitudinal cohort of patients undergoing carotid or femoral endarterectomy at two tertiary centers in the Netherlands. The study design has been described previously^28^. Transcriptomic data from 1,093 atherosclerotic plaque segments (AERNAS1, n=622; AERNAS2, n=471) together with the corresponding clinical and plaque phenotyping data were accessed through the AtheroPortal (Peters TS, Krebbers S, van der Laan SW, et al., medRxiv, 2025; accessed on june 2026 at <https://atheroportal.nl/app/>). A Kruskal-Wallis test was used to determine whether there are statistically significant differences between groups. Comparison between groups is performed using t-test for each group pair, and p-values are corrected for multiple testing using the Bonferroni correction.

***In vivo* experiments**

**Mouse model**

*Rev-erbα^-/-^* (*Nr1d1^-/-^*) mice were obtained from B. Vennström^29^ and backcrossed for 8 generations with wild-type mice on a sv129 background or for 3 generations with atherosclerosis-prone *Ldlr^-/-^* mice on a C57BL/6J background. Mice were housed in a 12-hour light/12-hour dark cycle. Male *Ldlr^-/-^ Rev-erbα^+/+^* and *Ldlr^-/-^Rev-erbα^-/-^* littermate mice were fed a standard chow diet *ad libitum* and sacrificed at 12 months (n=8 per genotype) and 18 months (n=10 per genotype) of age to investigate atherosclerosis and vascular calcification progression, respectively. During aging, 2 mice from the 18-months-old *Ldlr^-/-^Rev-erb^+/+^* group and 1 mouse from the 18-months-old *Ldlr^-/-^Rev-erb^-/-^* group died. Determination of mice number was based on literature^30,31^. No exclusion or inclusion criteria were used at this stage. No randomization was necessary here. Data analysis was performed in double-blinded manner. For isolation of primary aortic SMCs, thoracic aortas were isolated from 7-week-old sv129 *Rev-erbα^+/+^* and *Rev-erbα^-/-^* littermate mice fed a standard chow and further processed as described below.

**RNA extraction and quantitative PCR (qPCR)**

Total RNAs were isolated from descending aortic tissue and cells using Trizol reagent (Ambion, Life technology) and reverse transcribed into cDNA using the High-Capacity cDNA [reverse transcription](https://www-sciencedirect-com.proxy.insermbiblio.inist.fr/topics/medicine-and-dentistry/reverse-transcription) kit (4368813; Applied Biosystems, Waltham, USA). QPCR analysis was performed with the qPCR low Rox SYBR Master Mix dTTP blue (Takyon, Eurogentec) on an Mx3005P (Agilent Technologies), QuantStudio 3 or QuantStudio 6 Pro (Applied Biosystems) using specific primers (see Tables 1 and 2 for details). Relative quantification was performed using the *Ppia* gene encoding Cyclophilin A. Relative mRNA levels (2^-∆∆Ct^) were determined by comparing (a) the PCR cycle thresholds (Ct) for the gene of interest and the house keeping gene *Cyclophilin A* (∆Ct) and (b) ∆Ct values for treated/*Rev-erbα^-/-^* and control groups/*Rev-erbα^+/+^* (∆∆Ct).

**Microarray analysis**

Mice were housed in a 12-hour light/12-hour dark cycle. Male *Ldlr^-/-^ Rev-erbα^+/+^* and *Ldlr^-/-^Rev-erbα^-/-^* littermate mice were fed a standard chow diet *ad libitum* and sacrificed at 12 months (n=10 per genotype). During aging, 3 mice from the *Ldlr^-/-^Rev-erb^+/+^* group died. Total RNAs were isolated from descending aortas using Trizol reagent (15596018, Ambion, Life technology). The concentration and integrity were evaluated using the Bioanalyser 2100 (Agilent Biotechnologies). mRNAs (300 ng) were amplified, fragmented and labelled using the Gene Chip WT Plus Reagent kit (902280, Affymetrix, Santa Clara, USA). Labelled RNAs were hybridized to MoGene 2.0 ST gene chips (Affymetrix). Washing and final staining were performed on the Gene Chip Fluidics Station 450 (Affymetrix). DNA chips were scanned using the Gene Chip Scanner 3000 7G (Affymetrix). Microarray data were processed using Galaxy-based Interactive tools for ANalysis of data (GIANT) on a local instance of Galaxy^32^. [Gene set enrichment](https://www-sciencedirect-com.proxy.insermbiblio.inist.fr/topics/medicine-and-dentistry/gene-set-enrichment) analyses were performed using the GSEA tool (v.4.0.3) from Broad Institute. The 12,000 genes with the highest mean were used to perform Differential Expressed Gene, Fast Gene Set Enrichment Analysis (fGSEA) and interaction analysis in Phantasus^33^ (v1.31.1). DEG were assessed using Limma. A Principal Component Analysis (PCA) was performed on these set of 12,000 genes. For DEG and interaction analysis, a cutoff was set at 0.05 for the adjusted *P-value*. GO term over-representation analysis for biological pathways was performed on 1,266 upregulated genes in *Ldlr^-/-^Rev-erbα^-/-^* mice compared to littermate with an adjusted *P-value*<0.05 using DAVID ([https://davidbioinformatics.nih.gov](https://davidbioinformatics.nih.gov/)). FGSEA and interaction analysis were performed in Phantasus^33^ (v1.31.1). We calculated the rich factor defined as the ratio of differentially expressed gene number to the total gene number in each pathway.

**Single Cell RNA sequencing**

To analyze gene expression profiles at the single cell level, brachiocephalic arteries, aortic arches and thoracic aortas were perfused with RPMI 1640 medium (11875093, Gibco, Life Technologies) and harvested from *Ldlr^-/-^Rev-erbα^+/+^* and *Ldlr^-/-^Rev-erbα^-/-^* mice (n=5 per group). *Adventitia* was then removed from the vessels and atherosclerotic area were separated from heathy region using a stereo microscope (MZ6, Leica) and microdissection tools. Arteries were minced and incubated in a filtered enzyme cocktail (collagenase type CLS-4, 0.2 mg/mL, Worthingthon #WOLS04186; Dispase purified NPRO, 0.2 UI/mL, Worthington #WOLS02100; Lyophilized elastase ESL, 1 mg/mL, Worthington #WOLS02292; DNAse I DP, 0.2 mg/mL, Worthington #WOLS02139) for 45 minutes at 37°C. Cells were isolated from tissue debris using a 70 µm cell strainer and were centrifuged at 400 x*g* for 5 minutes at 4°C. Cells were then labelled with an anti-CD45-BV510 antibody (BLE103138, Biolegend) and with HTO antibodies to identify single samples (TotalSeq^TM^-A series from Hashtag 1 to Hashtag 8; Biolegends). Cells were sorted on a BD FACS ARIA III sorter to eliminate CD45^+^ cells and assess cell viability. Encapsulation was performed using the 10X Genomics Chromium Controller system with Chromium Next GEM Single Cell 3’ Reagents (Kit v3.1). Single cell cDNA and cell hashing HTO libraries were prepared according to manufacturer’s instructions (10X Genomics and Biolegend, respectively), and sequenced at Beijing Genomics Institute (BGI, Hong Kong). Final median read depth for gene expression was 30-70k reads/cell and 1500-1400 reads/cell for HTOs. Sequences were aligned to a reference genome (mm10-2020-A), obtained from 10X Genomics, using Cellranger (7.1.0) software using the option *--include-introns*. Cellranger output was initially processed using the Seurat pipeline (v4) for QC (mitochondrial transcripts < 5%, nFeature_RNA < 4500 and nCount_RNA > 200) and doublet removal (identified as ‘Singlet’ using DoubletFinder – ref: <https://www.cell.com/cell-systems/fulltext/S2405-4712(19)30073-0>) and HTO cell assignment was performed using *MultiSeqDemux* normalized across cells (margin=2) with default parameters. Libraries were integrated using STACAS in fully unsupervised mode^34^. Quality control and pre-processing led to the identification of 5,428 total CD45^-^ cells from atherosclerotic plaque including 2,029 nuclei from *Ldlr^-/-^Rev-erbα^+/+^* and 3,399 nuclei from *Ldlr^-/-^Rev-erbα^-/-^.* The Seurat object was converted to a cloupe file for further exploration in Loupe (v9.1.0, 10X Genomics). Unbiased analysis using the Seurat R package identified 14 cell clusters with a resolution of 0.8. Clusters (0.8 resolution) were annotated based on their differential gene expression calculated against all other cells. Positive marker genes for each cluster were analyzed using both EnrichR (<https://maayanlab.cloud/Enrichr/>) and CellMarker2.0 (<http://www.bio-bigdata.center/CellMarkerSearch.jsp>) softwares to annotate cell types (see table 3 for cell markers).

**Immunohistochemistry**

Hearts were perfused with PBS followed by 4% paraformaldehyde (PFA) and embedded in a cryogenic solution. Serial cryosections of 10 µm of the heart and aortic valves were obtained using a HM 560 (Microm Microtech). Sections were blocked with a Blocking Solution (SP-6000, Bloxall, Vector Laboratories) and a Protein Block solution (UltraVision, Thermo Scientific) and subsequently labelled with primary antibodies against MOMA-2 (sc59332, Santa Cruz) and CD4. Secondary antibodies (from Southern Biotech) conjugated to horseradish peroxidase protein polymer (MP-7404, NovaRED Immpress, Vector Laboratories) were revealed using a substrate solution (SK-4800, Vector RED, Vector Laboratories). Hearts sections were also stained with oil red O and alizarin red to visualize lipid and calcium deposition, respectively. Nucleus were stained with hematoxylin. Sections were scanned (9 images per sections, magnification 10X objective Nikon 10x/0.3 dry Plan Fluor) and quantified (28 sections per heart) using an inverted microscope (Nikon Eclipse Ti) and Nis Elements software. Alizarin red stained sections were scanned using the slide scanner Axioscan Z1 (Zeiss) and were quantified using ImageJ software (v2.16.0/1.54t).

**Immunofluorescence assays**

For confocal analysis, fixed ascending aorta and aortic sinus sections were incubated for 4 hours in a permeabilization/blocking solution (PBS1X, 5% donkey serum, 0.1% triton X-100). NLRP3 expression within macrophages was assessed using primary antibodies against NLRP3 (ab4207, Abcam, d1/200) and MOMA-2 (sc59332, Santa Cruz, d1/200). Images were acquired using an inverted LSM880 confocal laser-scanning microscope with a 10x objective (EC P1 N DIC 10x/0.3 dry, pixel size 0,354 x 0,354 µm) (Zeiss). Osteoblastic differentiation analysis was evaluated with a primary antibody against osterix (ab209484, Abcam, d1/500), and Cemip expression within atherosclerotic plaques was quantified using a primary antibody against Cemip (BS-8059R, Bio-Rad, d1/50). Antibodies against α-SMA (ab184675, Abcam, d1/200) and CD64 (MCA5997, Bio-Rad, d1/1000) were used to identify SMCs and macrophages, respectively. Sections were incubated overnight at 4°C with primary antibodies and 4 hours at room temperature with Alexa-coupled secondary antibodies (A-21208, A-21432, A-31573, Jackson laboratory, d1/500). Nuclei were stained with DAPI (Thermofischer Scientific). Incubation with second antibodies only were used as negative control to detect unspecific staining. Sections were scanned using the slide scanner Axioscan Z1 (Zeiss) and quantified using ImageJ software (v2.16.0/1.54t). Representative images acquired using an inverted LSM980 confocal laser-scanning microscope with a 20x objective (Plan-Apochromat 20x/0.8 dry, pixel size 0.127 x 0.127 µm) (Zeiss).

**Cell isolation and culture**

***Primary aortic Smooth Muscle Cells isolation and culture***

To isolate primary aortic SMCs, aortas were dissected from 7 to 12-week-old *Rev-erbα^+/+^* and *Rev-erbα^-/-^* male mice fed a standard chow. Aortas were cleared of the perivascular adipose tissue and adventitia and then digested to dissociate SMCs. These primary aortic SMCs were cultured in DMEM/F12 (11320033, Gibco, Life Technology), supplemented with GlutaMAX (1%, 35050061, Gibco, Life Technologies), gentamycin (0.4%, 15710, Gibco, Life Technologies) and 20% of fetal bovine serum (FBS) (FBS-12A, Capricorn Scientific, Clinisciences).

***MOVAS culture***

The Movas SMC line (ATCC, CRL-2797) was cultured in DMEM medium (11966-025, Gibco, Life Technologies) supplemented with 20% FBS (FBS-12A, Capricorn Scientific, Clinisciences), GlutaMAX (1%, 35050061, Gibco, Life Technologies), G418 (0.2%, A1720, Sigma Aldrich) and gentamycin (0.4%, 15710, Gibco, Life Technologies).

***In vitro* osteoblastic differentiation of SMC**

To obtain osteoblast-like cells, SMCs were cultured in DMEM (11966-025, Gibco, Life Technologies) supplemented with gentamycin (0.4%, 15710, Gibco, Life Technologies), Glutamax (1%, 35050061, Gibco, Life Technologies) and 10% of fetal bovine serum in presence of phosphate (Na_2_PO_4_・2H_2_O, 3 mM) (71643, Sigma-Aldrich) for 2 weeks. Osteoblast-like cells were then treated with recombinant murine IL-1β (0.1 ng/mL) (Biolegend) in osteoblast-like cell differentiation medium for one week.

**Calcium deposition Assay**

SMC were cultured in 24-well plates for 2 weeks, differentiated into osteoblastic cells and fixed in 4% PFA. Cells were stained with alizarin red and scanned (12 images per well, 3 wells per condition, 40X magnification, Objective Nikon ELWD 40x/0.6 dry S Plan Fluor). Calcium deposits were quantified using a Nikon Eclipse Ti microscope and Nis Elements software.

**TNAP activity Assay**

Proteins were extracted from osteoblast-like cells, and the alkaline phosphatase activity was assayed according to the supplier’s recommendations (SensoLyte pNPP Alkaline Phosphatase Assay Kit, AS-72146, Anaspec).

**Pharmacological treatments**

To activate Rev-erb, cells were treated with 6 µM hemin for 15 days (H9039, Sigma-Aldrich) during osteoblast-like cell differentiation.

**Chromatin Immunoprecipitation Assay**

ChIP assays were performed as described previously^35^  with the following modifications. SMCs were synchronized with 50% of horse serum for 2 hours. 39 hours post-synchronization, cells were cross-linked with 1% of formaldehyde solution (28908, ThermoScientific). Chromatin was sonicated during 35 cycles (30 seconds on, 30 seconds off). Precipitation was performed using the anti-Rev-erbα (1/50, #13418, Cell Signaling) and normal rabbit IgG (1/250, #2729, Cell Signalling Technologies) antibodies following manufacturer’s instructions. Rev-erbα-DNA complexes were immunoprecipitated, washed, eluted, purified and submitted to qPCR analysis. Crossing threshold (Ct) values were determined for each promoter and normalized to the Ct of input using the following equation: relative values = 2^-(Ct IP - Ct input)^. Data were expressed as a fold induction compared with the negative region using 2^-∆∆Ct^ with 2^-[(Ct IP region of interest – Ct input) – (Ct IP negative region – Ct input)]^. The oligonucleotides listed in Table 4 were used to amplify the negative region (a region of the genome with no transcriptional regulatory marks used to define the background) and the regions containing putative Rev-erbα response elements in the promoter of the *Sp7*, *Spp1* and *Cemip* genes. Identification of these putative RevRE sequences was performed using Promoter database. Enrichment in Rev-erbα binding was calculated using the comparative Ct method and expressed relative to control samples.

**Statistical Analysis**

Statistical analysis was performed on GraphPad Prism software (version 11.0.2) using either *t* test after the normality is verified or Mann-Whitney test for comparisons between 2 groups and one-way and Kruskal-Wallis or two-way ANOVA, after the normality is verified, with Šídák or *post-hoc* multiple comparisons of more than 2 groups, as indicated in the legends. An identification of statistical outliers was performed with a ROUT (Q=1%). Differences were considered significant when *P*<0.05. All *in vitro* experiments were repeated independently at least three times, and representative results are presented.

**Supplementary figures legend**

**Supplementary Figure 1: Alterations in circulating monocyte subsets** **in diabetic patients according to calcium score quartile and analysis of *REV-ERB* expression in different areas of atherosclerotic plaques.**

**A-D.** Quantification of leukocytes (A), monocytes (B), classical monocytes (C) and non-classical monocytes (D) based on their quartile classification according to CAC score in the AngioSafe 2 diabetic cohort. Data are shown as violin plots (Q1: n=218, Q2: n=220, Q3: n=128, Q4: n=102). Statistical significance was determined using the Kruskal-Wallis test, with * *p*<0.05 and ** *p*<0.01. **E.** Relative levels of *REV-ERBα* mRNA in CD14^+^CD16^-^ monocytes from diabetic patients with low CAC score (Q1 ; n=11) and high CAC score (Q4 ; n=11). Data are shown as min-to-max box, and statistical significance was assessed using an unpaired *t*-test, * *p*<0.05. **F.** Relative ***REV-ERBα*** mRNA expression in patients with type 2 diabetes with no or minor calcification (n=113) or moderate to heavy calcification (n=103), and in non-diabetic patients with no or minor calcification (n=398) or moderate to heavy calcification (n = 354) from the AtheroPortal cohort. Data are presented as min-to-max box plots. Statistical analysis was performed using a Kruskal Wallis test with *p-value*=0.038, then a *t*-test was performed, and *p-*value was corrected for multiple testing using the Bonferroni correction method (<https://atheroportal.nl/app/>), * *p*<0.05. **G.** Relative levels of *REV-ERBβ* mRNA in non-calcified (n=11) and calcified (n=22) areas of human endarterectomies from the OPAL cohort. Data are shown as min-to-max box, and statistical significance was assessed using an unpaired *t*-test, * *p*<0.05. **H.** Representative histological staining of human endarterectomy specimens used for laser capture microdissection to define distinct regions of atherosclerotic plaques. Lipid-rich and structural regions were identified by Oil Red O and H&E staining, vascularized regions by CD31/CD68 immunostaining, and calcified regions by Alizarin Red staining. Contrast and brightness adjustments were applied equally to each conditions using ImageJ2 software (2.16.0/1.5t).

**Supplementary Figure 2: Transcriptomic analysis of aortas from *Ldlr^-/-^* mice and List of human genes associated with vascular calcification used in transcriptomic analysis.**

**A-B.** Principal component analysis (PCA) (A) and heatmap (B) of differential gene expression in aortas of 18-month-old *Ldlr^-/-^Rev-erbα^+/+^* and *Ldlr^-/-^Rev-erbα^-/-^* mice. **C.** Bubble plot representation of untargeted GO biological pathways-based FGSEA analysis using whole aortic DEGs from 18-month-old *Ldlr^-/-^Rev-erbα^+/+^* and *Ldlr^-/-^Rev-erbα^-/-^* mice. **D.** Human vascular calcification gene used for analysis

**Supplementary Figure 3: Characterization of CD45^-^ cells in atherosclerotic lesions in mice.**

**A–B.** Heatmap showing the top 5 markers of each cell type (A) and violin plot showing the distribution of marker in each cluster (B) **C.** UMAP plots showing the expression of representative markers shown in panel B in the different clusters. Color-coding indicated expression of the indicated marker genes for each cluster. Color scale: purple, high expression; orange, low expression.

**Supplementary Figure 4: Cell frequency in *Ldlr^-/-^Rev-erbα^+/+^* and *Ldlr^-/-^ Rev-erbα^-/-^* atherosclerotic plaque.**

Histogram showing the frequency of the different cell types identified by single-cell RNA sequencing per genotype and per mouse. Statistical analysis was performed using a Mann-Whitney test, *p*-value are indicated.

**Supplementary Figure 5: Rev-erbα controls osteoblastic differentiation in SMCs by regulating transcription of osteoblastic markers.**

**A.** mRNA expression levels of the anti-calcifying osteoblastic markers *Mgp* (Matrice Gla Protein) in SMC isolated from *Rev-erbα^+/+^* mice treated or not with hemin (6 µM) and cultured with or without osteogenic differentiation medium (Na_2_PO_4_.2H_2_O, 3 mM) for 15 days. Data are mean ± SD (n=3 per condition). Statistical analysis was performed using a two-way ANOVA test followed by a Šídák *post-hoc* test to compare genotypes or treatments. **B.** Diagram illustrating putative Rev-erbα response elements (RevRE) in the *Spp1* promoter, obtained using MatInspector software. **C-H.** ChIP analysis of Rev-erbα occupancy of *Spp1* (C-G) and *Arntl* (Bmal1*)* (H) promoter response elements in synchronized primary aortic SMCs derived from *Rev-erbα^+/+^* and *Rev-erbα^-/-^* mice (n=3 per condition). Data represent means ± SEM. Statistical analysis was performed using a two-way ANOVA test followed by a Šídák *post-hoc* test. * *p*<0.05, ** *p*<0.01, **** *p*<0.0001.

**Supplementary Figure 6: Impact of *Rev-erbα* deficiency on NLRP3 inflammasome activation and IL-1β-induced osteoblastic differentiation in SMCs.**

**A.** Relative levels of the *NLRP3* inflammasome mRNA in atherogenic (n=41) versus healthy regions (n=42) of human endarterectomies the OPAL 1 cohort. Data are represented by a violin plot, analyzed by unpaired *t*-test, *** *p*<0.001. **B.** Expression levels of the *Nlrp3* inflammasome and the pro-inflammatory cytokines *Il-1β* and *Il-18* in aortas from 18-month-old *Ldlr^-/-^Rev-erbα^+/+^* and *Ldlr^-/-^Rev-erbα^-/-^* mice. Data are shown as means ± SEM (n=12 per genotype). Statistical analysis was performed using a Mann-Whitney test. *** *p*<0.05, *** *p*<0.001. **C-D.** Representative immunofluorescence staining of Nlrp3 in macrophages and lesional SMCs (C) from aortic sinus sections and their quantification (D) in aortas from 18-month-old *Ldlr^-/-^Rev-erbα^+/+^* and *Ldlr^-/-^Rev-erbα^-/-^* mice. Data are means ± SD (n=6-8 per genotype). Statistical analysis was performed using a Mann-Whitney test. *p<0.05. Contrast and brightness adjustments were applied equally to each condition using ImageJ2 software (2.16.0/1.5t). **E-F.** Representative staining (E) and quantification of calcium deposits (F) after alizarin red staining in SMCs isolated from *Rev-erbα^+/+^* and *Rev-erbα^-/^*^-^ mice cultured with or without osteogenic differentiation medium (Na_2_PO_4_•2H_2_O, 3 mM) ± IL-1β (0.1 ng/mL) for 7 days (12 fields per well, n=3 per condition). Data are means ± SD. Statistical analysis was performed using a two-way ANOVA test followed by a Šídák *post-hoc* test. **** *p*<0.0001.

**Supplementary Figure 7: Metabolic pathways and genes dysregulated by *Rev-erbα* deficiency during osteoblastic differentiation.**

**A-C.** Principal component analysis (PCA) plot (A), heatmap (B) and Venn diagram (C) showing gene expression profiles and the distribution of the number of genes specifically regulated in SMCs isolated from *Rev-erbα^+/+^* and *Rev-erbα^-/-^* mice, cultured with or without osteogenic differentiation medium (Na_2_PO_4_•2H_2_O, 3 mM) for 15 days (n= 3-5 per condition as indicated). **D.** FGSEA analysis identifying pathways whose gene expression is differentially regulated by Rev-erb-α during osteoblastic differentiation. **E**. ChIP analysis of Rev-erb-α occupancy of the *Arntl* (Bmal1) promoter in MOVAS cells, synchronized and cultured with or without osteogenic differentiation medium (Na_2_PO_4_•2H_2_O, 3 mM) (n=4 per condition). Data represent means ± SD. Statistical analysis was performed using a two-way ANOVA test followed by a Šídák *post-hoc* test. * *p*<0.05, ** *p*<0.01, *** *p*<0.001, **** *p*<0.0001. **F.** *Rev-erbα* and *Cemip* mRNA expression levels in SMC isolated from *Rev-erbα^+/+^* and *Rev-erbα^-/-^* mice, with or without *Cemip* knock-down, and cultured with or without osteogenic differentiation medium (Na_2_PO_4_•2H_2_O, 3 mM) ± IL-1β (0.1 ng/mL) for 48h (n=3 per condition). Data are shown as means ± SD. * *p*<0.05, ** *p*<0.01, **** *p*<0.0001.

**Supplementary Figure 8 : Graphical abstract**

Rev-erbα exerts pleitropic effects on vascular calcification, targeting multiple mechanistic pathways.

**Tables**

**Table 1: Mouse primers**

| **Gene** | **Sense primer sequence** | **Antisense primer sequence** |
| --- | --- | --- |
| *Ppia* | GCA TAC GGG TCC TGG CAT CTT GTC C | ATG GTG ATC TTC TTG CTG GTC TTG C |
| *Rev-erbα* | TGG CCT CAG GCT TCC ACT ATG | CCG TTG CTT CTC TCT CTT GGG |
| *Runx2* | CTT CAC AAA TCC TCC CCA AG | GAA TGC GCC CTA AAT CAC TG |
| *Sp7* | CCT CTC GAC CCG ACT GCA GAT C | AGC TGC AAG CTC TCT GTA ACC ATG AC |
| *Bglap* | TAG TGA ACA GAC TCC GGC GCT A | TGT AGG CGG TCT TCA AGC CAT |
| *Spp1* | TCC CTC GAT GTC ATC CCT GT | CCC TTT CCG TTG TTG TCC TG |
| *Tnfrsf11b* | AGT CCG TGA AGC AGG AGT | CCA TCT GGA CAT TTT TTG CAA A |
| *MGP* | GCA ACC CTG TGC TAC GAA TC | TGC GTT CCT GGA CTC TCT TT |
| *Cemip* | TCC CAC TTG CTT ACG CTC AG | AAC AGC TTC CTC CCT CAG GT |
| *Tnap* | ATC GAC GTG ATC ATG GGT GG | GGT CAA TCC TGC CTC CTT CC |
| *Nlrp3* | CCC TTG GAG ACA CAG GAC TC | GAG GT G CA GTT GTC TAA TTC C |
| *IL-1β* | GCC ACC TTT TGA CAG TGA TGA G | CCT GAA GCT CTT GTT GAT GTG C |
| *IL-18* | CTG GGG TTC ACT GGC ACT TTG ATC | TTG GCC GAC TTC ACT GTA CAA CCG |

**Table 2: Human primers**

| **Gene** | **Sense primer sequence** | **Antisense primer sequence** |
| --- | --- | --- |
| *PPIA* | GCA TAC GGG TCC TGG CAT CTT GTC C | ATG GTG ATC TTC TTG CTG GTC TTG C |
| *REV-ERBα* | GAG TCA AGG TCC AGT TTG AAT GAC | CAG CTT GGT GAT GTT GCT GGT GCT C |
| *REV-ERBβ* | AAT CGA GTG CAC CTG GGA TG | CCG AAA GAA ACC CTT ACA GCC |

**Table 3. Cellular populations and markers identified in aortic atherosclerotic lesions**

| **Cell type / phenotype** | **Gene markers used for identification** | **References** |
| --- | --- | --- |
| Foam cells / macrophage-like foam cells | *Plin2, Abcg1, Cd68, Ptprc* (CD45)*, Acta2* (αSMA)*, CD11b* | ^3^ |
| Endothelial cells | *Cdh5* (VE-cadherin)*, Pecam1* (CD31) | ^45^ |
| Lymphatic endothelial cells | *Cdh5 (VE-cadherin), LYVE1* | ^45^ |
| Vascular endothelial cells | *Cdh5 (VE-cadherin), End1* | ^45^ |
| Erythrocytes | *Hba-a1, Alas2* | ^46^ |
| Early erythrocytes | *Hba-a1, Alas2, Car2* | ^46^ |
| Contractile SMC | *Acta2, Cnn1, Myh11* | ^13,47,48^ |
| Proliferative SMCs | *Acta2, Ccnd1, Tff2* | ^13,50,51^ |
| Migrating SMCs associated with the fibrous cap | *Acta2, Myh11, Phactr1* | ^11,13^ |
| Osteochondroblast progenitors / osteogenic cells | *Sox9* | ^47,52^ |
| Differentiated osteoblasts / osteogenic cells | *Sox9, Postn* | ^47,52,53^ |
| Pericyte-derived osteoblast-like cells | *Sox9, Postn, Wif1* | ^47,52–54^ |
| Chondroblast cells | *Sox9, Spp1, Acan, Tnfsr11b, Col2a1* | ^47,52,53,55^ |
| Osteoprogenitor cells | *Sox9, Anxa8, Comp, Tmem100* | ^47,52,56,63^ |
| Fibroblast-lineage / fibro-osteoblast cells | *Sox9, F3, ApoD, C7, Gdf10, Smoc2* | ^47,52,53,57–60^ |
| Chondrofibromyocytes | *Sox9, Ptx3, Tmem100, Has1* | ^47,52,61–63^ |
| Osteoblasts | *Sox9, Comp, Cxcl14* | ^47,52,56^ |

**Table 4: Mouse primers used for chromatin immunoprecipitation experiments.**

| **Gene** | **Sense primer sequence** | **Antisense primer sequence** |
| --- | --- | --- |
| Sp7 Prom +301 | ACT TCC AGG GTC CTC CTC AT | TCT TTG ACT GCC CTC TGA CC |
| Spp1 Prom -6520 | CCA CAG ACT CCC TTT CCA GG | CCC ATG TGC TGG AAA CTA TGC |
| Spp1 Prom -3070 | CAG TGA AAC CTT TCT GTG CC | GAA GGT GGT CTT ACA TGA G |
| Spp1 Prom -2120 | GGC ATC AAC AGA GCG GGA C | GCA TCC TTA GAG GCC GAC TG |
| Spp1 Prom -560 | CTT TGT TTC GTT TTG TTG TGA GG | GCT CTG TTT TGT ACT CCG G |
| Cemip Prom -2819 | CGT CTT TCT CTG TGG TGG TG | GTT TTG TTG AAG GCC AAG GA |
| Cemip Prom +350 | AAG CTA TGG GGA GGA GAG GA | GAT AGG CAG GCA GGC AGA C |
